# Metformin inhibits nuclear egress of chromatin fragments in senescence and aging

**DOI:** 10.1101/2024.12.16.628669

**Authors:** Takuya Kumazawa, Yanxin Xu, Tara C. O’Brien, Ji-Won Lee, Yu Wang, Murat Cetinbas, Ruslan I. Sadreyev, Nabeel Bardeesy, Chia-Wei Cheng, Bin He, Zhixun Dou

**Author notes:** Correspondence: Zhixun Dou.

## Abstract

Chronic inflammation is a hallmark of aging and contributes to many age-associated diseases. Metabolic intervention is a strategy to modulate inflammation. However, the connection between inflammation and metabolism during aging remains poorly understood. A mechanism driving chronic inflammation involves cytoplasmic chromatin fragments (CCFs), which appear in senescent cells and aged tissues, activating the cGAS-STING pathway. The size of the CCFs exceeds the capacity of the nuclear pore complex, raising the question of how chromatin fragments enter the cytoplasm. Here, we report that chromatin fragments exit the nucleus via nuclear egress, a membrane trafficking process at the nuclear envelope that shuttles large complexes from the nucleus to the cytoplasm. Inactivating critical nuclear egress ESCRT-III or Torsin proteins results in accumulation of chromatin fragments at the nuclear membrane, thereby impairing the activation of cGAS-STING and senescence-associated inflammation. Notably, nuclear egress of CCFs is inhibited by glucose limitation or metformin treatment. This is due to AMPK phosphorylation and autophagic degradation of the ESCRT-III component, ALIX. Metformin treatment in naturally aged mice downregulates ALIX protein and blocks cGAS activation and chronic inflammation in the small intestine. Together, our study defines a central mechanism linking nutrient sensing and chronic inflammation, two distinct hallmarks of aging, and suggests a new approach to suppress age-associated inflammation by targeting the nuclear egress of chromatin fragments.

## Main

While acute inflammation restrains infection, chronic inflammation in the absence of infection promotes aging and age-associated diseases ^1^. Targeting age-associated chronic inflammation without impairing beneficial effects of acute inflammation is therefore an important biomedical objective. Metabolic interventions and chronic inflammation are closely connected: caloric restriction, rapamycin, and metformin have all been reported to suppress chronic inflammation and promote healthspan and/or lifespan in animal models ^2^. However, these metabolic interventions have side effects and/or are difficult to implement. Understanding the relationship between metabolism and inflammation could enable the development of more specific therapies for age-associated inflammatory diseases.

One mechanism that drives chronic inflammation involves cytoplasmic chromatin fragment (CCF)^3^. CCFs are observed in senescent cells, naturally aged tissues, and in diseased tissues associated with chronic inflammation ^4–6^. CCFs are sensed by cGAS, leading to activation of the cGAS-STING pathway ^5,7,8^. We and others have shown that CCF-cGAS-STING signaling contributes to senescence and age-associated chronic inflammation ^5,7–9^. However, the mechanisms underlying CCF formation remain poorly understood, posing challenges for developing targeted therapies. In particular, we have limited knowledge of how chromatin fragments in the nucleus enter the cytoplasm to become CCFs. Live-cell imaging experiments revealed that CCFs are generated via nuclear membrane blebbing, which eventually releases chromatin fragments into the cytoplasm ^4,10^, indicating the involvement of a membrane trafficking process.

A clue is provided by studies of viral nuclear egress. Several DNA viruses, such as herpesviruses, replicate and package their genomes into viral capsids in the host cell nuclei. Because these viral capsids exceed the size limit of the nuclear pore complex, they are transported to the cytoplasm through nuclear egress ^11–13^. This involves disruption of the nuclear lamina, passage through the inner nuclear membrane, and budding out of the outer nuclear membrane ^11,12,14^. In addition to viral nuclear egress, cells use nuclear egress to transport large ribonucleoprotein particles (RNPs) from the nucleus to the cytoplasm ^12,15^. Viral nuclear egress and CCF nucleus-to-cytoplasm transport share several similarities: (1) both viral capsids and CCFs are large DNA-protein complexes that exceed the capacity of the nuclear pore complex, (2) both processes involve nuclear lamina remodeling, and (3) both involve a transient nuclear membrane blebbing step. These parallels led us to hypothesize that chromatin fragments exiting the nucleus are mediated by nuclear egress.

In this study, we demonstrate that the nucleus-to-cytoplasm transport of chromatin fragments is mediated by nuclear egress in senescence and aging. We further report that AMPK and metformin suppress the nuclear egress of CCFs. Importantly, inhibition of CCF nuclear egress does not impair acute inflammation triggered by cytosolic DNA, suggesting a new approach for specifically targeting age-associated chronic inflammation without compromising acute inflammation.

## Results

### ESCRT-III complex mediates CCF nucleus-to-cytoplasm transport

We began our study by assessing the size of CCF in cellular senescence, a stable form of cell cycle arrest associated with inflammation (detailed conditions to induce and validate senescence are described in Methods). The appearance of CCFs coincides with the induction of the senescence-associated secretory phenotype (SASP), which occurs in the late stage of senescence after the cell cycle arrest has been established ^4–6^. CCFs in senescent primary human IMR90 fibroblasts were found to have an average diameter of ∼2 μm (Fig. 1a), which is about 20 times larger than the capacity of the nuclear pore complex (∼100 nm). Similar sizes of CCFs were observed under various senescence-inducing conditions, including ionizing irradiation (IR), the DNA damaging agent etoposide, or activated Ras (HRasV12) (Fig. 1a).

**Figure 1.**
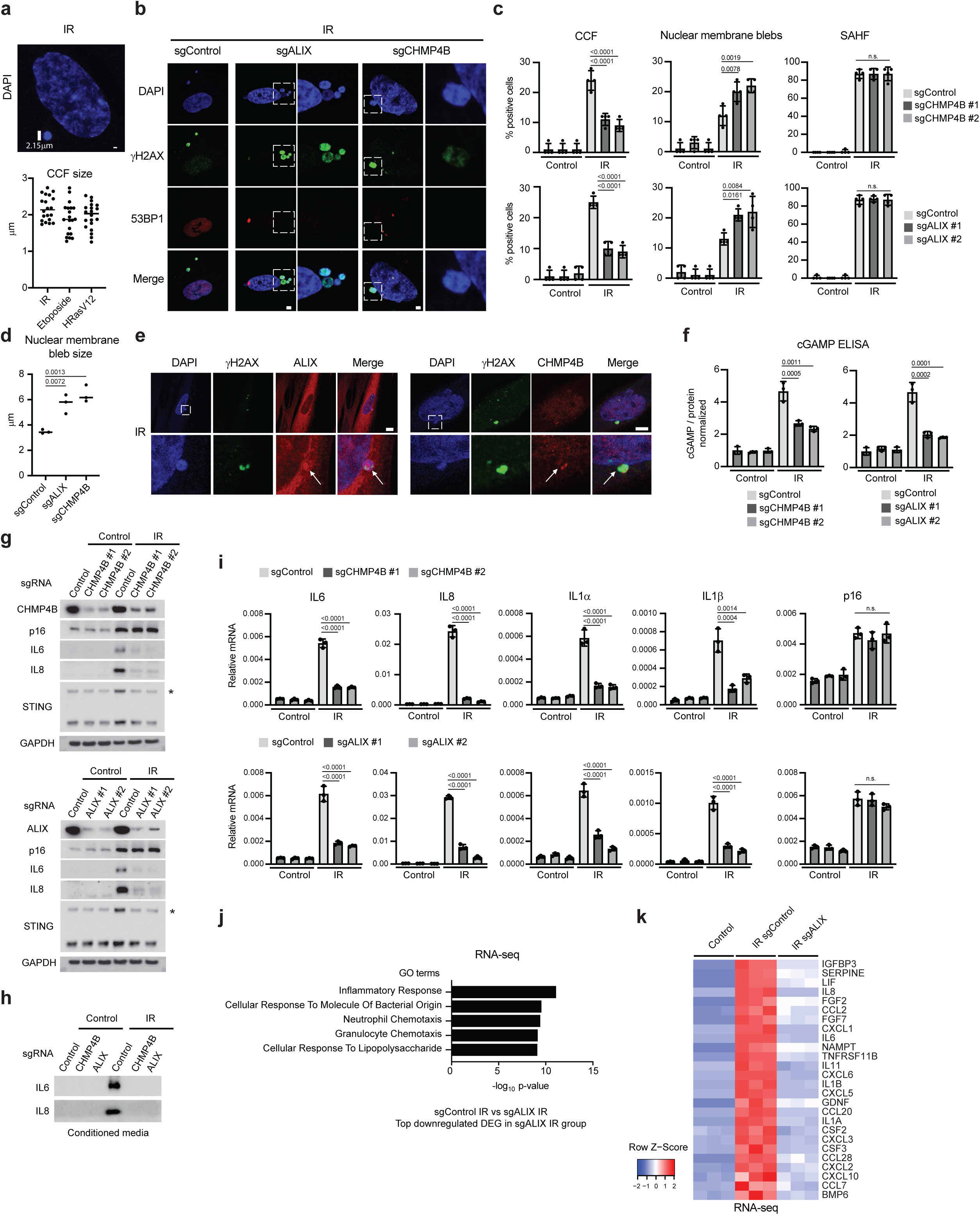
ESCRT-III complex is required for CCF formation in senescence. **a**, CCF size measurements in senescent IMR90 cells. Image shown is senescence induced by IR for 14 days with CCF size labeled (top). Scale bar: 1 μm. Quantification of various forms of senescence is shown (bottom). The major axis of the oval shape is quantified. **b**, IMR90 cells were stably infected with CRISPR KO constructs encoding sgControl, sgALIX, or sgCHMP4B. The cells were then induced to senescence by IR (harvested on day 14), followed by antibody staining and imaging under a confocal microscopy. Note that the KO cells display less CCF and abnormal nuclear membrane blebs. Scale bar: 1 μm. **c**, Related to **b**. Quantification of CCF in the cytoplasm, nuclear membrane blebs, and senescence-associated heterochromatin foci (SAHF). Data presented are the mean values with s.d. from four randomly selected fields with over 200 cells. P values are calculated with one-way ANOVA coupled with Tukey’s post hoc test and labeled on the graphs. n.s.: non-significant. **d**, Quantification of the length of nuclear membrane blebs, with mean values labeled. See Materials and Methods for details. Results are from three independent experiments. P values are calculated with one-way ANOVA coupled with Tukey’s post hoc test. **e**, Senescent IMR90 cells induced by IR were stained with ALIX and CHMP4B antibodies and imaged under a confocal microscopy. Note the presence of ALIX and CHMP4B on nuclear membrane blebs. Scale bar: 5 μm. **f**, cGAMP ELISA measurements of cGAS activation in IMR90 proliferating or IR-induced senescent cells. Results were normalized to total protein and were presented as mean values with s.e.m.. P values are calculated with one-way ANOVA coupled with Tukey’s post hoc test. **g**, IMR90 cells with indicated genotypes were induced to senescence by IR and analyzed by immunoblotting. STING westerns were performed under non-reducing conditions. * denotes STING dimer. **h**, The conditioned media were analyzed by immunoblotting. The amounts of media loaded were normalized to total cellular proteins. See Materials and Methods for details. **i**, RT-qPCR analyses for CHMP4B or ALIX KO cells. The expression of genes were normalized to those of Lamin A/C. Results shown are the mean values with s.d.. P values are calculated with one-way ANOVA coupled with Tukey’s post hoc test. n.s.: non-significant. **j** and **k**, RNA-seq analyses of sgControl and sgALIX senescent IMR90 cells, performed in three replicates. Gene Ontology (GO) analysis of top downregulated genes in sgALIX senescent cells (**j**). Heatmap presentation of key SASP genes from three replicates (**k**).

To investigate how chromatin fragments are transported from the nucleus to the cytoplasm, we examined proteins involved in viral nuclear egress. The host ESCRT-III complex has been reported to mediate scission of the inner nuclear membrane, enabling viral capsids to enter the perinuclear space (the space between inner and outer nuclear membranes) ^12^. Subsequently, the host Torsin complex enables the budding of the viral capsids from the outer nuclear membrane ^14^.

To inactivate the ESCRT-III complex, we disrupted CHMP4B, which mediates membrane scission, and ALIX, a regulatory component required for ESCRT-III activity. CRISPR-mediated knockout of CHMP4B or ALIX, using two independent sgRNAs for each gene, resulted in impaired cytoplasmic localization of CCFs in IR-induced senescent cells, accompanied by accumulation of chromatin fragments at the nuclear membrane blebs (Fig. 1b and quantified in Fig. 1c; also see Extended Data Fig. 1a for additional examples). CCFs stain positive for γH2AX but negative for 53BP1, which typically colocalizes with γH2AX in the nucleus (Ref^4^ and Fig. 1b). The accumulation of γH2AX-positive, 53BP1-negative chromatin fragments at the nuclear membrane blebs indicates a failure for CCF budding off the nuclei. Indeed, nuclear membrane blebs in CHMP4B- or ALIX-deficient cells exhibited increased frequency and size (Fig. 1b and quantified in Fig. 1c and 1d). By contrast, senescence-associated heterochromatin foci (SAHF), a marker for chromatin reorganization ^16^, were unaffected by CHMP4B or ALIX deficiency (Fig. 1b and quantified in Fig. 1c). Consistent with the roles of CHMP4B and ALIX in CCF trafficking, both proteins were detected at the nuclear membrane blebs in senescent cells (Fig. 1e; also see Extended Data Fig. 1b for additional examples), revealed by antibody staining of endogenous proteins (the specificities of the antibodies are confirmed using CRISPR KO cells, showing the absence of staining, in Extended Data Fig. 1c). CHMP4B and ALIX are not enriched at the CCFs after CCFs reach the cytoplasm (Extended Data Fig. 1d), indicating that the ESCRT-III complex acts specifically during the nuclear membrane blebbing step.

Next, we examined the consequences of reduced CCF cytoplasmic localization in ESCRT-III-deficient senescent cells. Notably, these cells exhibited impaired activation of the cGAS-STING pathway, as assessed by cGAMP ELISA (Fig. 1f) and STING dimerization (Fig. 1g; STING dimer is marked by *). Consequently, ESCRT-III-deficient cells exhibited impaired induction of the SASP, as measured at the protein (Fig. 1g) and secreted levels (Fig. 1h), in accordance with the effects on associated mRNAs (Fig. 1i). In contrast, the induction of p16^ink4a^ (hereafter “p16”) and senescence-associated beta-galactosidase (SA-β-gal) were unaffected (Fig. 1g, 1i, and Extended Data Fig. 1e). The reduction of the SASP was also confirmed by RNA-seq. Analysis of differentially expressed genes (DEGs) revealed that ESCRT-III-deficient cells have impaired expression of a broad panel of pro-inflammatory genes (Extended Data Fig. 1f), enriched for key features of the SASP (Fig. 1j), also presented in a heatmap (Fig. 1k).

To evaluate the generalizability of these findings, we examined various senescence-inducing conditions and cell strains beyond IR-induced senescence in IMR90. The dependence of CCF transport and SASP activation on ESCRT-III was confirmed under etoposide-induced senescence in IMR90 cells (Extended Data Fig. 2), IR-induced senescence in BJ cells (Extended Data Fig. 3, top), and etoposide-induced senescence in A549 cells (Extended Data Fig. 3, bottom). In addition to senescence, we investigated acute cytosolic dsDNA responses by transfecting cells with interferon stimulatory DNA (ISD) and harvested the cells at 24 hours, and found that the induction of pro-inflammatory genes was largely unaffected in ESCRT-III-deficient cells (Extended Data Fig. 4). This indicates that ESCRT-III does not directly regulate the cGAS-STING pathway. In sum, our results collectively suggest that CCF nucleus-to-cytoplasm transport is regulated by ESCRT-III-mediated trafficking, which is essential for activating the cGAS-STING-SASP pathway in senescence.

### Nuclear ESCRT-III regulates CCF exit from the nucleus

A key regulator of ESCRT-III activity is VPS4, an AAA-ATPase that facilitates the exchange of factors in and out of the ESCRT-III complex and recycles ESCRT-III proteins for repeated action^17^. We evaluated the impact of a dominant negative (DN) VPS4 mutant compared to wild-type (WT) VPS4 in senescence. The VPS4-DN mutant significantly reduced CCF in the cytoplasm, accumulated nuclear membrane blebs (Extended Data Fig. 5a), and inhibited the expression of SASP genes (Extended Data Fig. 5b). In contrast, VPS4-DN had minimal effects on SAHF formation (Extended Data Fig. 5a), p16 expression (Extended Data Fig. 5b), and acute responses to dsDNA transfection (Extended Data Fig. 5c), indicating that VPS4 acts on CCFs to regulate the SASP.

The ESCRT-III complex is known to regulate membrane trafficking at various subcellular locations, including the plasma membrane, cytoplasm, and nuclear envelope ^18,19^. To investigate whether the nuclear pool of ESCRT-III regulates CCF, we added a 3x nuclear localization signal (NLS) to VPS4, driving its localization to the nucleus (Extended Data Fig. 5d). Expression of the 3xNLS-tagged VPS4-DN mutant blocked the appearance of CCF in the cytoplasm, accumulated nuclear membrane blebs (Extended Data Fig. 5e), and suppressed SASP gene expression (Extended Data Fig. 5f). These results highlight that the nuclear pool of ESCRT-III is essential for CCF nucleus-to-cytoplasm shuttling in senescence.

### Torsin proteins mediate CCF nucleus-to-cytoplasm trafficking

We next investigated the role of Torsin complex, which is required for the budding of viral capsids and large RNP from the outer nuclear membrane ^14,15^. Torsin is an AAA-ATPase that localizes at the perinuclear space and the lumen of the endoplasmic reticulum (ER) ^20^. We found that inactivation of Torsin 1A (TOR1A), a key component of the Torsin complex, resulted in the accumulation of nuclear membrane blebs and impaired appearance of CCF in the cytoplasm (Fig. 2a-2c; Extended Data Fig. 6a). Consequently, cGAS activation, as measured by cGAMP ELISA, was reduced in TOR1A-deficient senescent cells (Fig. 2d). This was accompanied by a reduced activation of the SASP, measured by immunoblotting (Fig. 2e), RT-qPCR (Fig. 2f), and RNA-seq (Fig. 2g). The induction of SAHF, p16, and SA-β-gal was unaffected by TOR1A deficiency (Fig. 2b and 2f, Extended Data Fig. 6b). Torsin 1B (TOR1B) is another member of the Torsin complex, which shares functional redundancy with TOR1A ^20^. Disruption of TOR1B also blocked CCF and the SASP, although double KO of TOR1A and TOR1B generated more pronounced effects (Extended Data Fig. 6c and 6d). Similar to ESCRT-III, Torsin is required for CCF formation and the SASP across multiple cell strains and under various senescence-inducing conditions (Extended Data Fig. 7). As an important control, inactivation of Torsin had minimal impact on acute pro-inflammatory gene expression triggered by dsDNA transfection (Extended Data Fig. 8). Collectively, these findings identify Torsin proteins as critical mediators of CCF nuclear egress in senescence.

**Figure 2.**
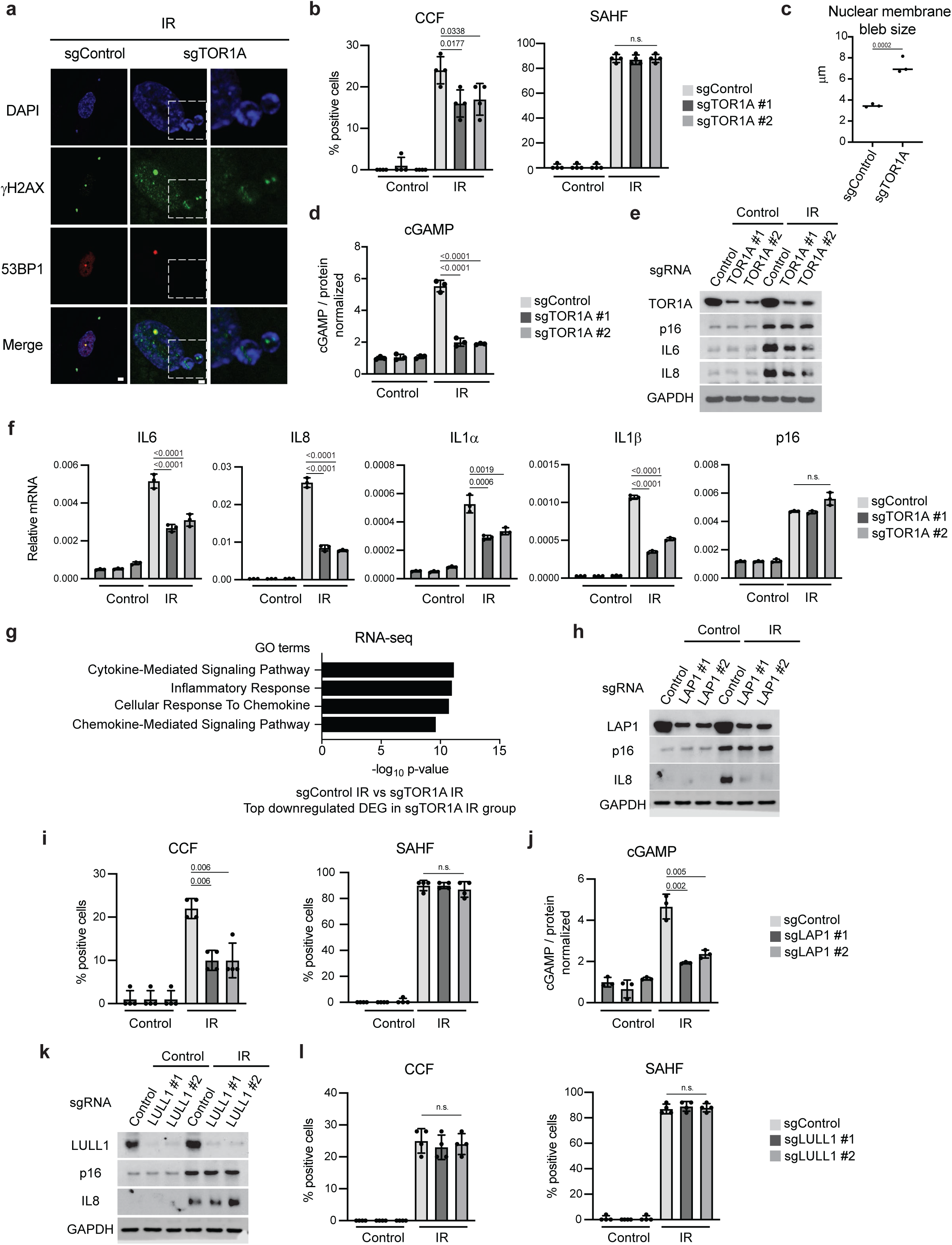
Torsin is required for nuclear egress of CCF. **a**, IMR90 cells were stably infected with CRISPR KO vector encoding sgControl or sgTOR1A, followed by induction of senescence with IR. The cells were fixed, stained, and imaged as indicated. Scale bar: 5 μm. **b**, Related to **a**. Quantification of CCF in the cytoplasm and senescence-associated heterochromatin foci (SAHF). Data presented are the mean values with s.d. from four randomly selected fields with over 200 cells. P values are calculated with one-way ANOVA coupled with Tukey’s post hoc test. n.s.: non-significant. **c**, Quantification of the length of nuclear membrane blebs, with mean values labeled. See Materials and Methods for details. Results are from three independent experiments. P values are calculated with one-way ANOVA coupled with Tukey’s post hoc test. The same sgControl group was used as in Fig. 1d and hence one-way ANOVA was used to compare multiple groups. **d**, cGAMP ELISA measurement of cGAS activation in IMR90 proliferating or IR-induced senescent cells. Results were normalized to total protein and were presented as mean values with s.e.m.. P values are calculated with one-way ANOVA coupled with Tukey’s post hoc test. **e**, IMR90 cells with indicated genotypes were induced to senescence by IR and analyzed by immunoblotting. **f**, RT-qPCR analyses for control and TOR1A KO cells. The expression of genes were normalized to that of Lamin A/C. Results shown are the mean values with s.d.. P values are calculated with one-way ANOVA coupled with Tukey’s post hoc test. **g**, RNA-seq analyses of sgControl and sgTOR1A senescent IMR90 cells, performed in three replicates. Gene Ontology (GO) analysis of top downregulated genes in sgALIX senescent cells is shown. **h**-**j**, IMR90 cells were stably infected with CRISPR KO vector encoding sgControl or sgLAP1. The cells were then induced to senescence by IR and analyzed by immunoblotting (**h**), imaging (**i**), or cGAMP ELISA (**j**). Bar graphs present mean values with s.d. for imaging analyses from four randomly selected fields with over 200 cells. Bar graph for cGAMP ELISA shows mean values with s.e.m. normalized to total proteins. P values are calculated with one-way ANOVA coupled with Tukey’s post hoc test. n.s.: non-significant. **k** and **l**, IMR90 cells were stably infected with CRISPR KO vector encoding sgControl or sgLULL1. The cells were then induced to senescence by IR and analyzed by immunoblotting (**k**) or imaging (**l**). Bar graphs present mean values with s.d. from four randomly selected fields with over 200 cells. P values are calculated with one-way ANOVA coupled with Tukey’s post hoc test. n.s.: non-significant.

Torsin proteins are present in both the perinuclear space and the ER lumen, raising the question of which pool of Torsin is necessary for CCF nuclear egress. To address this, we examined the binding partners of Torsin. The catalytic activity of Torsin is stimulated by interactions with LAP1 and LULL1. On its own, Torsin exhibits minimal catalytic activity ^21^. LAP1 is localized at the nuclear lamina, stimulating Torsin activity at the perinuclear space, while LULL1 is localized at the cytoplasmic side of the ER, stimulating membrane remodeling of the ER ^21–23^. We thus disrupted LAP1 or LULL1 by CRISPR-mediated KO, and found that LAP1, but not LULL1, is required to induce the CCF-cGAS-SASP pathway in senescence (Fig. 2h-2l). These results suggest that the nuclear membrane pool of Torsin is critical for driving the nuclear egress of CCF.

### Glucose limitation downregulates ALIX and inhibits CCF nuclear egress

Dietary and metabolic interventions are being investigated as strategies to mitigate chronic inflammation during aging ^2,24^. While CCF plays a key role in mediating chronic inflammation, its connection to metabolic interventions remains unclear. To explore this relationship, we investigated whether nutrient limitation affects proteins involved in CCF nuclear egress. Notably, glucose limitation dramatically downregulated ALIX protein levels (Fig. 3a), whereas amino acid or serum deprivation for the same duration failed to do so (Fig. 3a). Other proteins involved in CCF nuclear egress were not downregulated by these treatments (Extended Data Fig. 9a). The reduction of ALIX upon glucose limitation was observed in both control cells and senescent cells induced by IR or etoposide (Fig. 3a), suggesting that ALIX regulation by glucose availability is a general mechanism.

**Figure 3.**
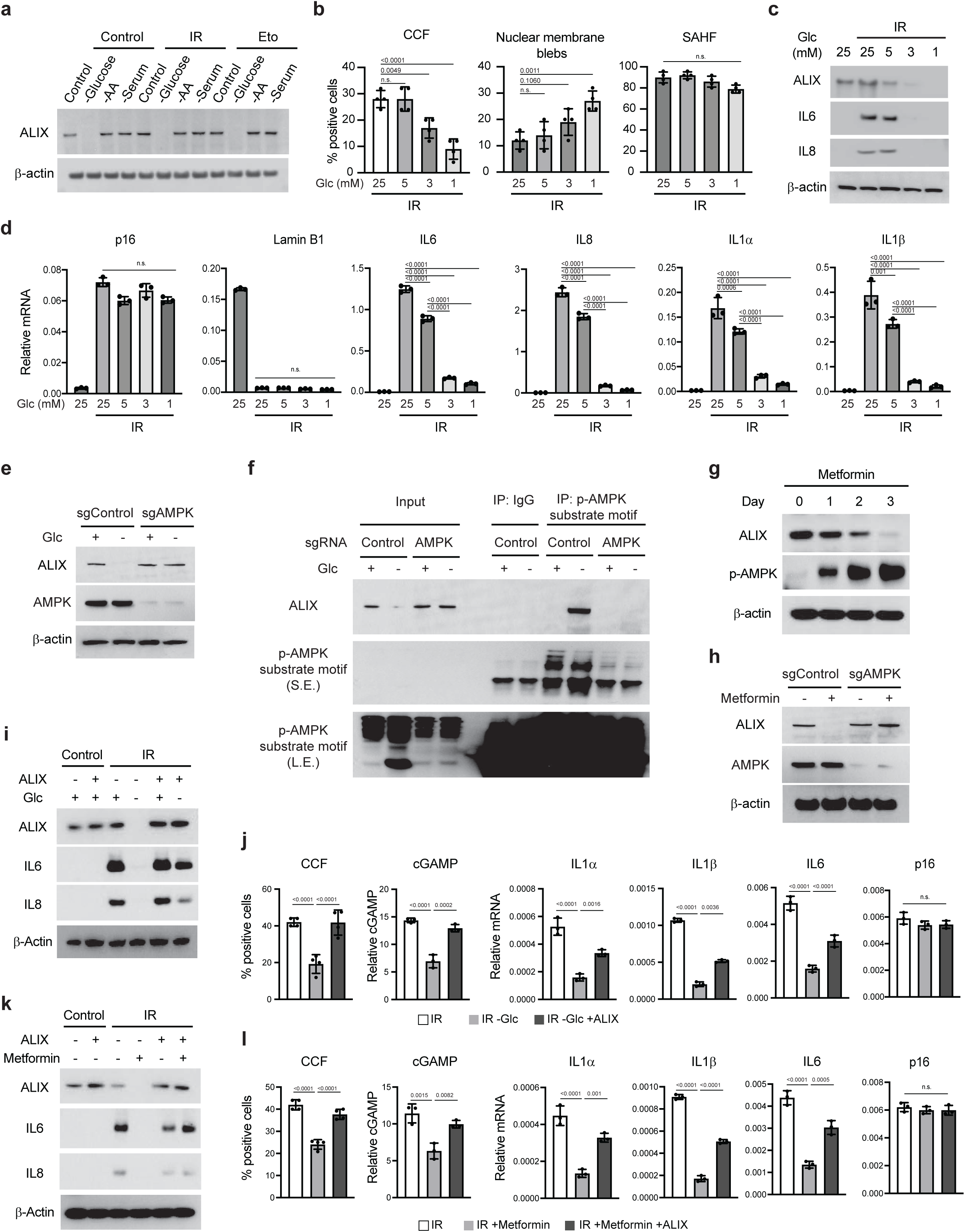
Glucose limitation and metformin inhibit CCF and the SASP. **a**, IMR90 cells under proliferating or senescent conditions were left untreated or treated with nutrient limitation for 4 days. Senescence was induced as described in Materials and Methods and nutrient limitation was performed for the last 4 days. These treatments caused minimal cell death. The cell lysates were analyzed by immunoblotting. **b**-**d**, IMR90 cells were induced to senescence by IR for 8 days, followed by culturing in media harborbing various glucose concentrations for 6 days. The cells were then analyzed by imaging (**b**), immunoblotting (**c**), or RT-qPCR (**d**). Bar graphs in (**b**) present mean values with s.d. from four randomly selected fields with over 200 cells. Bar graphs in (**d**) present mean values with s.d.. P values are calculated with one-way ANOVA coupled with Tukey’s post hoc test. n.s.: non-significant. **e**, IMR90 cells stably expressing CRISPR KO vector encoding sgControl or sgAMPKα were left untreated or cultured in no glucose media for 3 days. The lysates were analyzed by immunoblotting. **f**, Cells as in **e** were lysed in 1% SDS buffer followed by 95 ℃ boiling. SDS was then diluted to 0.1%, followed by immunoprecipitation with p-AMPK substrate motif antibody and immunoblotting. This denature IP condition ensures that protein-protein interactions were disrupted and thus the ALIX brought down was a direct consequence of AMPK phosphorylation. **g**, IMR90 cells were cultured in 5 mM media in the presence of 5 mM metformin for indicated days and were analyzed by immunoblotting. **h**, Cells with indicated genotypes were cultured in 5 mM glucose media and treated with 5 mM metformin for 3 days, followed by immunoblotting. **i-l**, IMR90 cells were stably expressed with lentiviral vector or lentiviral vector expressing ALIX. The cells were then induced to senescence by IR for 8 days, followed by culturing in 1 mM glucose media for 6 days (**i** and **j**) or treated with 5 mM metformin for 6 days in 5 mM glucose media (**k** and **l**). The cells were then analyzed by immunoblotting (**i** and **k**), imaging for cytoplasmic CCF, ELISA for cGAMP, or RT-qPCR for SASP gene expression (**j** and **l**). Bar graphs showing CCF present mean values with s.d. from four randomly selected fields with over 200 cells. Bar graphs showing cGAMP ELISA and RT-qPCR present mean values with s.e.m. (cGAMP) or s.d. (RT-qPCR). P values are calculated with one-way ANOVA coupled with Tukey’s post hoc test.

Given the essential role of ALIX in facilitating CCF nuclear egress and the SASP (Fig. 1 and Extended Data Fig. 1), we next examined whether glucose availability regulates CCF. Glucose concentrations in culture media were manipulated as follows: DMEM with high glucose (25 mM) served as the standard, while concentrations below the physiological level of 5 mM were considered glucose-limited. Cells were induced to senescence by IR for 8 days in 25 mM glucose media, followed by culturing in media with various glucose concentrations for additional 6 days. This procedure revealed that glucose limitation dose-dependently reduced CCF in the cytoplasm while increasing nuclear membrane blebs (Fig. 3b). This was accompanied by a reduction of SASP gene expression observed at the protein (Fig. 3c) and the mRNA levels (Fig. 3d). The induction of p16 and SAHF as well as loss of Lamin B1 were largely unaltered by glucose limitation in senescent cells (Fig. 3b and 3d). Notably, the effects of glucose limitation on CCF and the SASP phenocopied those observed in ALIX KO cells (Fig. 1), suggesting that CCF nuclear egress is regulated by glucose availability.

### Metformin/AMPK inhibits CCF generation via ALIX downregulation

AMPK is a central sensor of energy stress and is activated under low glucose conditions ^25^, prompting us to investigate its potential involvement in ALIX regulation. CRISPR-mediated KO of AMPK abrogated ALIX downregulation upon glucose limitation (Fig. 3e). Furthermore, using a phospho-AMPK substrate motif antibody, which specifically recognizes the Rxx(pS/pT) motif characteristic of AMPK substrates ^26^, we observed that ALIX was recognized by this antibody under denaturing immunoprecipitation conditions. This recognition occurred specifically upon glucose deprivation and in an AMPK-dependent manner (Fig. 3f).

Having established a critical role of AMPK in mediating ALIX downregulation, we tested AMPK-activating compounds. Both metformin and AICAR treatment under physiological glucose conditions (5 mM) reduced ALIX protein levels (Fig. 3g and Extended Data Fig. 9b). The reduction in ALIX by metformin was abolished in AMPK KO cells (Fig. 3h), indicating that metformin downregulates ALIX via AMPK activation.

Next, we evaluated the connection between glucose limitation/metformin, ALIX, and the CCF-cGAS-SASP pathway. Glucose limitation reduced ALIX protein levels (Fig. 3i), CCF, and cGAMP production (Fig. 3j), which can be rescued by forced ALIX expression (Fig. 3i and 3j). Furthermore, while glucose limitation reduced the SASP gene expression, forced expression of ALIX partially restored this defect (Fig. 3i and 3j). Metformin treatment also blocked the CCF-cGAS-SASP pathway in a manner that was significantly rescued by ectopic expression of ALIX (Fig. 3k and 3l). These data collectively suggest that ALIX is a key mediator linking glucose limitation or metformin to the suppression of the SASP.

To place metformin’s effects in a broader context, we investigated its impact on other cytosolic DNA mechanisms beyond CCF. Metformin had minimal impact on acute dsDNA transfection-induced pro-inflammatory gene expression (Extended Data Fig. 9c), indicating that metformin does not directly target cGAS or STING. In addition, we tested metformin’s effect on mitochondrial DNA (mtDNA) release into the cytosol, which has been reported to promote the cGAS-STING-SASP pathway in senescence and aging ^27^. Metformin had minimal impact on mtDNA release into the cytosol in senescent cells (Extended Data Fig. 9d). These results suggest that CCF, rather than cGAS-STING or mtDNA, is a major target of metformin in senescence.

### ALIX is degraded by autophagy upon glucose limitation

We subsequently investigated the mechanism underlying glucose limitation-induced ALIX downregulation. ALIX mRNA levels were unaffected by nutrient deprivation (Fig. 4a), indicating a post-transcriptional regulation. AMPK can promote protein degradation by macroautophagy (hereafter referred to as “autophagy”). Inhibiting autophagy by using an ULK1/2 inhibitor or Atg7 deletion suppressed ALIX reduction upon glucose limitation (Fig. 4b and 4c). Furthermore, ALIX showed reduced intensity and formed punctate structures colocalizing with the lysosomal marker LAMP1 under glucose starvation (Fig. 4d). To further visualize ALIX autophagic degradation, we generated an mCherry-GFP-ALIX construct, in which an orange signal (due to merged mCherry and GFP) indicates that the fusion protein is in a neutral pH environment, whereas a red signal (due to acidic quenching of GFP) indicates that the fusion protein is in acidic lysosomes ^28^. Under glucose limitation, mCherry-GFP-ALIX exhibited cytoplasmic puncta, including red-only puncta, which colocalized with LC3B and/or LAMP1 (Extended Data Fig. 10), confirming that ALIX undergoes autophagy-lysosome degradation upon glucose limitation.

**Figure 4.**
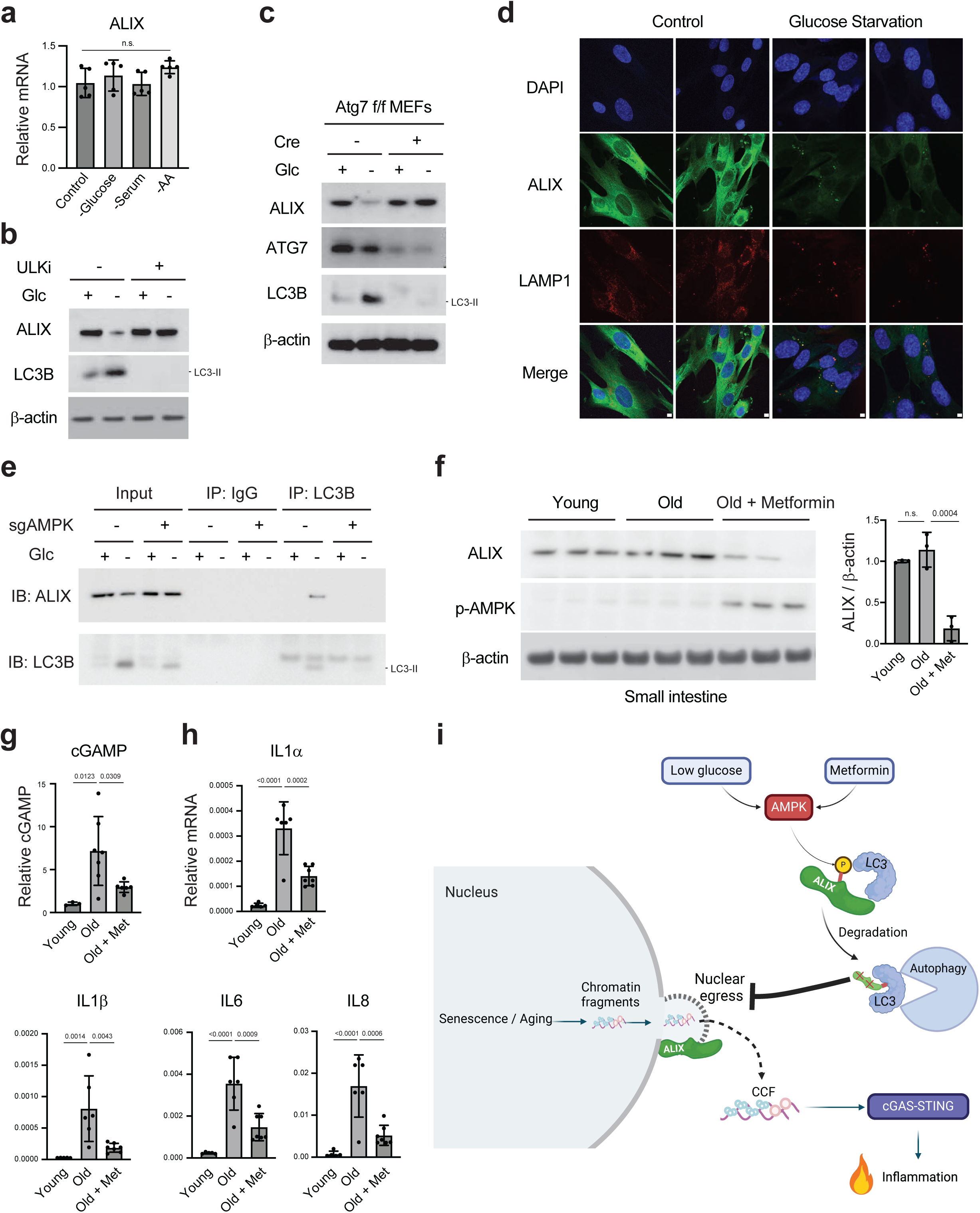
Autophagy mediates ALIX degradation and roles of metformin in the mouse intestine. **a**, Related to Fig. 3a. Proliferating IMR90 cells were treated with nutrient deprivation followed by RT-qPCR analyses. Results shown are mean values with s.d., normalized to Lamin A/C. n.s.: non-significant. P values are calculated with one-way ANOVA coupled with Tukey’s post hoc test. **b**, IMR90 cells were treated with 5 μM ULK1/2 inhibitors and were cultured in glucose-free media supplemented with 1 mM sodium pyruvate for 2 days. The lysates were analyzed by immunoblotting. **c**, MEFs with indicated genotypes were left untreated or cultured in glucose-free media supplemented with 1 mM sodium pyruvate for 3 days. The lysates were analyzed by immunoblotting. **d**, IMR90 cells were left untreated or cultured in 1 mM glucose media for 36 hours. The cells were then fixed, stained with antibodies as indicated, and imaged under a confocal microscopy. Scale bar: 5 μm. **e**, IMR90 cells with indicated genotypes were subjected to glucose limitation for 1 day, followed by immunoprecipitation with LC3B antibody and immunoblotting analyses. **f**, Immunoblotting of mouse small intestines from young (<4 mo), old (>24 mo), and old treated with metformin. **g**, cGAMP ELISA was performed using small intestines from young, old, and old treated with metformin. Results shown are mean values with s.e.m.. P values are calculated with one-way ANOVA coupled with Tukey’s post hoc test. **h**, RT-qPCR analyses of intestines from indicated groups. Results shown are mean values with s.e.m., normalized to those of Lamin A/C. P values are calculated with one-way ANOVA coupled with Tukey’s post hoc test. **i**, Schematic illustration of nuclear egress and its regulation by glucose and metformin. During senescence and aging, chromatin fragments can be produced in the nucleus and shuttled to the cytoplasm via nuclear egress, mediated by ESCRT-III (including ALIX) and Torsin proteins. CCF in the cytoplasm activates the cGAS-STING pathway, driving chronic inflammation. Low glucose levels or metformin activates AMPK, which phosphorylates ALIX, promoting ALIX binding to the autophagy adaptor LC3 and ALIX autophagic degradation. The loss of ALIX impairs nuclear egress of CCF, thereby inhibiting cGAS-STING activation and inflammation.

The degradation of selective autophagy substrates can be mediated by interaction with autophagy adaptor protein LC3, often in a phosphorylation-dependent manner ^29^. We found that LC3B immunoprecipitated ALIX at endogenous levels upon glucose limitation, in an AMPK-dependent manner (Fig. 4e). In summary, our findings reveal that AMPK mediates ALIX autophagic degradation upon glucose limitation, thereby suppressing the CCF-cGAS-STING pathway and the SASP.

### Metformin inhibits cGAS-STING in the intestine of aged mice

Lastly, we investigated the role of our newly identified pathway in naturally aged mice. While metformin exerts pleiotropic effects on aging in mice and monkeys ^30–33^, the mechanisms by which metformin inhibits chronic inflammation remain poorly understood. Given that the intestine is a major target of metformin ^34,35^, we focused on this organ to study its effects on the cGAS-STING pathway.

Naturally aged mice (over two years old) were fed *ad libitum* and provided with metformin in drinking water for three weeks. This treatment resulted in a substantial reduction of ALIX protein levels in the small intestine of metformin-treated groups (Fig. 4f). While cGAMP levels were increased in the intestines of aged mice, metformin treatment significantly reduced this increase (Fig. 4g), consistent with our in vitro findings (Fig. 3). In line with the impaired cGAS activation, the expression of key inflammatory cytokines was substantially reduced in the metformin-treated group (Fig. 4h). These results indicate that the cytosolic DNA sensing cGAS-STING pathway is a major target of metformin in the small intestine of naturally aged mice.

## Discussion

In this study, we demonstrate that CCFs undergo nuclear egress for nucleus-to-cytoplasm shuttling. This discovery helps fill a major gap in our understanding of the mechanisms underlying CCF generation, a critical event driving chronic inflammation in senescence and aging. We further reveal that CCF nuclear egress can be suppressed by glucose limitation or metformin via activation of AMPK. This finding establishes a direct link between nutrient metabolism and chronic inflammation (illustrated in Fig. 4i). Notably, blocking CCF nuclear egress does not impair acute cytosolic DNA-induced inflammatory responses, unlike inhibitors of cGAS, STING, or NFκB, which broadly suppress both acute and chronic inflammation. Therefore, targeting CCF nuclear egress represents a promising therapeutic strategy for treating age-associated chronic inflammatory diseases.

Previous studies have reported several mechanisms for metformin’s anti-aging effects, including activation of AMPK and autophagy, inhibition of mTOR, DNA damage, and NFκB ^30,36^. Our study identifies a new target for metformin – CCF. We showed that metformin downregulates ALIX, thereby inhibiting CCF nuclear egress and cGAS activation in senescence and aging. Restoration of ALIX largely reversed the effects of metformin on CCF and cGAS activation and partially rescued SASP gene expression (Fig. 3). The incomplete rescue of SASP gene expression by ALIX restoration suggests that additional factors are involved. This aligns with existing evidence that gene expression is an energy-demanding process, and AMPK may also inhibit the SASP through mTOR and NFκB ^37^. It is important to note that ALIX regulates other membrane trafficking processes beyond nuclear egress ^17–19^, cautioning potential side effects of metformin in the intestine.

Our study raises important questions for future research. First, the connection between chromatin fragments in the nucleus and the nuclear egress complexes remains unclear. HSV-1 nuclear egress involves viral UL31 and UL34 proteins that recruit CHMP4B and ALIX to the nuclear periphery, with ESCRT-III essential for viral nuclear egress ^12^. The mechanisms driving ESCRT-III to nuclear membrane blebs in senescent cells are yet to be addressed. Second, the factors mediating nuclear membrane bleb formation remain to be elucidated. Cytoskeletal remodeling of the nuclear membrane is likely involved and deserves future exploration. Lastly, beyond the context of senescence, we envision a new function for metformin in blocking herpesviruses, due to its activity to downregulate ALIX required for viral nuclear egress ^12^. The risk of herpesvirus reactivation increases with age, leading to complications such as shingles ^38,39^, which warrants further studies of metformin in aging.

## Acknowledgements

This project was conceived via discussion with Eric Baehrecke. We thank Maria Grazia Vizioli for piloting the conditions for this project and members of the Dou lab and Peter D. Adams lab for technical assistance and discussions. We acknowledge the microscopy core facility of Center for Regenerative Medicine at Massachusetts General Hospital for assistance on confocal microscopy, and the next-gen sequencing core at Massachusetts General Hospital for assistance on RNA-seq. This project is supported by NIH R01AG082785 (Z.D. and C.-W.C), NIH R35GM137889, UH3CA268117, R21AG073894, Hevolution/AFAR New Investigator Award, and Glenn Foundation for Medical Research and AFAR Grant for Junior Faculty (Z.D.), as well as NIH R01AI148148 (B.H.). YX is supported by Glenn Foundation for Medical Research Postdoctoral Fellowships in Aging Research from American Federation for Aging Research (AFAR).

## Author contributions

B.H. and Z.D. conceived the project. T.K. conducted most of the experiments unless stated. Y.X. contributed part of imaging results and data analyses. T.O. and Y.W. contributed part of the RNA-seq data. J.-W.L. contributed mtDNA analysis. M.C. and R.I.S. contributed computational analyses of RNA-seq. N.E.-B. supervised AMPK experiments. C.-W.C. supervised intestine experiments. B.H. supervised nuclear egress experiments. Z.D. supervised the study and provided most of the funding support. Z.D. wrote the paper. All authors discussed the manuscript.

The authors declare no competing financial interests.

## Methods

### Cell culture and treatment

All cells were cultured in Dulbecco’s Modified Eagle Medium (DMEM) high glucose (Thermo Fisher Scientific) supplemented with 10% fetal bovine serum (FBS) (Thermo Fisher Scientific), 100 units/mL penicillin, and 100 μg/mL streptomycin (Thermo Fisher Scientific), unless indicated otherwise. The cells were intermittently cultured with plasmocin (Invivogen) and were confirmed to be negative for mycoplasma using MycoAlert PLUS Mycoplasma Detection Kit (Lonza). Primary IMR90, BJ, and wild-type or Atg7 f/f MEFs were described previously ^5,10,40^. IMR90 were authenticated by next-gen sequencing ^10^. Primary cells were cultured in physiological oxygen (3%) incubators. A549 cells were acquired from American Type Culture Collection (ATCC).

For glucose limitation, cells were cultured in DMEM no glucose media (Thermo Fisher Scientific), supplemented with 10% dialyzed FBS (Thermo Fisher Scientific), 100 units/mL penicillin, and 100 μg/mL streptomycin (Thermo Fisher Scientific). For media with various glucose concentrations, glucose solution (Thermo Fisher Scientific) was added to glucose-free media. Sodium pyruvate (1 mM, Thermo Fisher Scientific) was added to 0 mM or 1 mM glucose media to enhance cell survival during glucose limitation. For experiments involving 5 mM glucose, DMEM low-glucose (Thermo Fisher Scientific) was used. Metformin (Selleckchem, 5 mM final concentration) treatments were done in DMEM low-glucose media. AICAR (APEXBIO, 1 mM final concentration) was also used for AMPK activation. ULK-101 (Selleckchem, 5 μM final concentration) was used for ULK1/2 inhibition. ISD (interferon stimulatory DNA) was from InvivoGen and was transfected using Lipofectamine 2000 (Thermo Fisher Scientific). Senescence-associated beta-galactosidase (SA-β-gal) was performed using a cellular senescence assay kit (Millipore Sigma) according to manufacturer’s instructions.

### Senescence induction and validation

IMR90 cells were used within population doublings of 40. The strain reaches replicative senescence around population doubling of 80. For etoposide-induced senescence, IMR90 cells at approximately 60-70% confluency were treated with 50 μM etoposide for 48 hours and harvested on day 10 to day 12. For ionizing radiation (IR), cells were irradiated with X-Rad 320 (Precision X-Ray Irradiator) at 20 Gy and harvested on day 14. For HRasV12-induced senescence, a retroviral vector encoding HRasV12 was used to infect cells. Following selection, cells were cultured for 7 more days. To establish senescence in BJ cells, cells were irradiated with X-Rad 320 (Precision X-Ray Irradiator) at 20 Gy and harvested on day 12. A549 was treated with 5 μM etoposide in cultured media and harvested on day 14. These conditions reproducibly induced senescence, confirmed by cell cycle arrest measured by the lack of Cyclin A and phosphorylated Rb, EdU less than 5% positivity, and SA-β-gal over 90% positivity, as described in the current and in our previous studies ^5,6,10,40–42^.

### Mice experiments

All mice experiments were performed in compliance with the Institutional Animal Care and Use Committee at Massachusetts General Hospital (Protocol # 2018N000182). Young (<4 mo) and aged (>24 mo) C57BL/6 mice were acquired from National Institute of Aging (NIA). Both male and female mice are included. Metformin (Selleckchem) was provided to mice in drinking water (0.35 mg/mL). Mice were fed *ad libitum* during metformin treatment, and were euthanized for tissue harvesting after 3 weeks of metformin treatment.

### Antibodies

Primary antibodies used include: gH2AX (Cell Signaling Technology 80312S), 53BP1 (Abcam ab21083), ALIX (for IF: Abcam ab275377, for WB: Santa Cruz sc-53540), CHMP4B (for IF: Abcam ab105767, Aviva systems biology ARP53340_P050, for WB: Invitrogen PA5-100092), p16 (BD Biosciences G175-405), IL6 (Cell Signaling Technology 12153), IL8 (Abcam ab18672), STING (Cell Signaling Technology 13647), GAPDH (Cell Signaling Technology 5174S), TOR1A (Thermo MA5-15094, Abcam ab34540), LAP1 (Abcam ab86307), LULL1 (Thermo 24769-1-AP), β-actin (Thermo PA5-143810), AMPKα (Cell Signaling Technology 5831), Phospho-AMPKα T172 (Cell Signaling Technology 2535), Phospho-AMPK substrate motif (Cell Signaling Technology 5759), IgG (Cell Signaling Technology 3900), LC3 (for IP and WB: Cell Signaling Technology 83506, for IF: Cell Signaling Technology 3868S), ATG7 (Cell Signaling Technology 8558S), LAMP1 (Developmental Studies Hybridoma Bank H4A3), HA (Sigma H3663), Flag (Sigma F-1804), VPS4A (Santa Cruz sc-393428).

Secondary antibodies used for western blotting include: Goat anti-Mouse IgG, IRDye® 680RD (Licor 926-68070), Goat anti-Mouse IgG, IRDye® 800RD (Licor 926-32210), Goat anti-Rabbit IgG, IRDye® 680CW (Licor 926-68071), Goat anti-Rabbit IgG, IRDye® 800CW (Licor 926-32211), Goat Anti-Mouse IgG (H + L)-HRP Conjugate (Bio Rad 1706516), Goat Anti-Rabbit IgG (H + L)-HRP Conjugate (Bio Rad 1706515), and Mouse Anti-Rabbit IgG (Light-Chain Specific) (Cell Signaling Technology 93702), Mouse Anti-Rabbit IgG Conformation Specific (Cell Signaling Technology 5127S).

Secondary antibodies used for immunofluorescence (IF) include: Goat anti-Mouse IgG Alexa Fluor 488 (Thermo Fisher Scientific A-11001), Goat anti-Mouse IgG Alexa Fluor 555 (Thermo Fisher Scientific A21422), Goat anti-Mouse IgG Alexa Fluor 647 (Thermo Fisher Scientific A-21236), Goat antiRabbit IgG, Alexa Fluor 488 (Thermo Fisher Scientific R-37116), Goat anti-Rabbit IgG Alexa Fluor 555 (Thermo Fisher Scientific A-21428), Goat anti-Rabbit IgG Alexa Fluor 647 (Thermo Fisher Scientific A-21446).

### Western blot

Western blotting was performed as described previously ^40,42^. Cells or tissues were lysed in a buffer containing 20 mM Tris, pH 7.5, 137 mM NaCl, 1 mM MgCl_2_, 1 mM CaCl_2_, 1% NP-40, supplemented with 1×Halt protease and phosphatase inhibitor cocktail (Thermo Fisher Scientific) and benzonase (Novagen) at 12.5 U/mL. The lysates were rotated at 4 °C for 30 minutes and boiled at 95 °C in the presence of 1% SDS. The resulting supernatants were subjected to electrophoresis using NuPAGE Bis-Tris precast gels (Thermo Fisher Scientific).

After transferring to nitrocellulose membrane, 5% milk in TBS was used to block the membrane at room temperature for 1 hour. Primary antibodies were diluted in 5% BSA in TBS supplemented with 0.1% Tween 20 (TBST) and incubated at 4 °C overnight. The membrane was washed 3 times with TBST, each for 10 minutes, followed by incubation of secondary antibodies at room temperature for 1 h, in 5% milk in TBST. The membrane was washed again 3 times and imaged by film or by an Odyssey imager (Licor Odyssey CLx 2000). STING dimer western blots were performed without adding a reducing agent when boiling the samples with loading dye.

For western blotting of conditioned media, control or senescent cells were cultured in corresponding media for 2 to 3 days before use. The media were then collected and filtered with a 0.45-μm polyvinylidene difluoride (PVDF) filter (Millipore) to remove cells and debris. The resulting supernatant was used for immunoblotting. The amounts of media used for immunoblotting were quantified based on the protein concentrations of cell lysates, and media corresponding to equal amounts of total cellular proteins were loaded into protein gels.

### Immunofluorescence and quantification

Immunofluorescence was performed as described previously ^40,42^. Cells were fixed in 4% paraformaldehyde in PBS for 30 minutes at room temperature. Cells were washed twice with PBS, and permeabilized with 0.5% Triton X-100 in PBS for 10 minutes. After washing twice with PBS, cells were blocked with 10% BSA in PBS for 1 hour at room temperature. Cells were then incubated with primary antibodies in 5% BSA in PBS supplemented with 0.1% Tween 20 (PBST) overnight at 4 °C. The next day, the cells were washed four times with PBST, each for 10 minutes, followed by incubation with Alexa Fluor-conjugated secondary antibody (Thermo Fisher Scientific), in 5% BSA/PBST for 1 hour at room temperature. The cells were then washed four times with PBST, incubated with 1 μg/mL DAPI (Thermo Fisher Scientific) in PBS for 10 minutes, and washed twice with PBS. The slides were mounted with ProLong Diamond (Thermo Fisher Scientific) and imaged with a Leica TCS SP8 fluorescent confocal microscope. Quantification of % positive cells was done under identical microscopy settings between samples. Over 200 cells from 4 randomly selected fields were analyzed.

The sizes of CCF and nuclear membrane blebs were quantified under Leica confocal software. For CCF, the major axis of the oval shape was presented as CCF size. For nuclear membrane blebs, the distance between the distal tip of the bleb and the nearest nuclear membrane was quantified.

### Immunoprecipitation

Cells were lysed in IP buffer containing 20 mM Tris, pH 7.5, 137 mM NaCl, 1 mM MgCl_2_, 1 mM CaCl_2_, 1% NP-40, supplemented with 1×Halt protease and phosphatase inhibitor cocktail (Thermo Fisher Scientific) and benzonase (Novagen) at 12.5 U/mL. The lysates were rotated at 4 °C for 30 minutes. The supernatant was then incubated with antibody-conjugated Dynabeads (Thermo Fisher Scientific) and rotated at 4°C overnight. The beads were then washed 4 times, collected by magnet stand, and boiled with 1×NuPAGE loading dye. Samples were analyzed by western blotting.

For immunoprecipitation involving phospho-AMPK substrate motif antibody, a denaturing condition was used. After lysing the cells as mentioned above, 1% SDS (final concentration) was added to the lysates, followed by 95°C boiling for 10 min. SDS was then diluted to 0.1% using the lysis buffer, followed by immunoprecipitation with the phospho-AMPK substrate motif antibody. This denaturing condition ensures that protein-protein interactions are disrupted, and thus the signal detected is a direct result of ALIX phosphorylation.

### cGAMP ELISA

Cells or tissues were lysed freshly using M-PER Mammalian Protein Extraction Reagent (Thermo) supplemented with 1×Halt protease and phosphatase inhibitor cocktail (Thermo Fisher Scientific). cGAMP ELISA was performed using 2’3’-cGAMP ELISA Kit (Cayman). Data are presented as cGAMP levels normalized to total proteins.

### Plasmids and viruses

pLentiCRISPRv2 was used to clone CRISPR constructs. Control CRISPR was described previously ^40,41^. The following sequences were used:

CHMP4B sg #1: CACCGAATGCCAACACCAACACCG, AAAACCGGTGTTGGTGTTGGCATTC

CHMP4B sg #2: CACCGGGCGGCCCGACCCCCCAGG, AAACCCTGGGGGGTCGGGCCGCCC

ALIX sg #1: CACCGCAGGCCCAGTACTGCCGCG, AAACCGCGGCAGTACTGGGCCTGC

ALIX sg #2: CACCGCAGGCCCAGTACTGCCGCG, AAACCGCGGCAGTACTGGGCCTGC

TOR1A sg #1: CACCGAGCATGTGGAAAGTGCAATG, AAACCATTGCACTTTCCACATGCTC

TOR1A sg #2: CACCGCGAGTGGAAATGCAGTCCCG, AAACCGGGACTGCATTTCCACTCGC

TOR1B sg #1: CACCGCACATGCACTCACATTACCG, AAACCGGTAATGTGAGTGCATGTGC

TOR1B sg #2: CACCGCAGAGTCGATACAAACAGG, AAACCCTGTTTGTATCGACTCTGC

LAP1 sg #1: CACCGACCTTCGCTCTCGACCACGG, AAACCCGTGGTCGAGAGCGAAGGTC

LAP1 sg #2: CACCGACAGTGTCTTACGTAACCCA, AAACTGGGTTACGTAAGACACTGTC

LULL1 sg #1: CACCGTATTTGTTGAGAAGTCATGG, AAACCCATGACTTCTCAACAAATAC

LULL1 sg #1:CACCGAAGGGTCAAAACCATCTCAG, AAACCTGAGATGGTTTTGACCCTTC

AMPKα1 sg #1: CACCGCTGGTGTGGATTATTGTCAC, AAACGACCACACCTAATAACAGTGC

AMPKα1 sg #2: CACCGGTAGATATATGGAGCAGTG, AAACCACTGCTCCATATATCTACC

ALIX as well as VPS4 wild-type and mutants’ cDNA were generated by GenScript, cloned into pGenLenti vector, and verified by Sanger sequencing. mCherry-GFP vector was described previously ^40^. VPS4 DN (E228Q) was used.

Lentiviral and retroviral vectors were packaged as described previously ^5,10,40^.

### RT-qPCR

RT-qPCR was described previously ^40,42^. Briefly, mRNA was extracted from cells or tissues using RNeasy Mini Kit (Qiagen), with a DNase I (Qiagen) digestion step to minimize genomic DNA contamination. Reverse transcription (RT) was done using High-Capacity RNA-to-cDNA Kit (Thermo), and then quantitative PCR (qPCR) was performed using a qPCR machine (BioRad CFX 384 Real-Time System). Results were normalized to Lamin A/C.

The following primers were used for RT-qPCR of human cells:

Lamin B1: CTCGTCGCATGCTGACAGAC, GATCCCTTATTTCCGCCATCT

p16: CCAACGCACCGAATAGTTACG, CCATCATCATGACCTGGATCG

IL6: CACCGGGAACGAAAGAGAAG, TCATAGCTGGGCTCCTGGAG

p21: CTCAGGGGAGCAGGCTGAAG, AGAAGATCAGCCGGCGTTTG

IL8: ACATGACTTCCAAGCTGGCC, CAAATCAGGAAGGCTGCC

IL1α: AGTGCTGCTGAAGGAGATGCCTGA, CCCCTGCCAAGCACACCCAGTA

IL1β: CTCTCTCCTTTCAGGGCCAA, GAGAGGCCTGGCTCAACAAA

Lamin A/C: AGCTGAAAGCGCGCAATACC, GGCCTCCTTGGAGTTCAGCA

ALIX: CATGCAGGGCAGTGAGGTTG, CCATCTTGAGCCAGGGCTGT

ISG15: AGGCGCAGATCACCCAGAAG, TTTGTCCACCACCAGCAGGA

ISG54: GAGCATGCTGACCAGGCAGA, CTGAGAGTCGGCCCATGTGA

ISG56: ATGGGCAGACTGGCAGAAGC, AGCAAGGCCCATCCTTCCTC.

The following primers were used for RT-qPCR of mouse cells/tissues:

IL-1α: TTCAAGGAGAGCCGGGTGAC, TGCTGATCTGGGTTGGATGG

IL1β: TCGCAGCAGCACATCAACAA, GCTGCCACAGCTTCTCCACA

IL6: CCGTGTGGTTACATCTACCCT, CGTGGTTCTGTTGATGACAGTG

CXCL1: CCATGGCTGGGATTCACCTC, CCAAGGGAGCTTCAGGGTCA

GAPDH: TGCATCCTGCACCACCAACT, ACGCCACAGCTTTCCAGAGG

Lamin A/C: TTCCCTGGAGACCGAGAACG, CACCTCTCGGCTGACCACCT.

### Cytosolic mitochondrial DNA quantification

A published protocol was used ^27^ with slight modifications. The cell pellets were lysed in digitonin buffer (150 mM NaCl, 50 mM HEPES pH 7.4, and 25 μg/ml digitonin (Sigma-Aldrich, D141)) for 10 min at room temperature. The digitonin-permeabilized cells were centrifuged at 150×g at 4 °C for 10 min. The resulting supernatant was further centrifuged at 20,000×g for 10 min to obtain the cytosolic fraction in the supernatant. This cytosolic fraction was then purified using a QIAquick Nucleotide Removal kit (Qiagen, 28306). In addition to the cytosolic fraction, whole-cell DNA was extracted, using a DNeasy Blood & Tissue kit (Qiagen, 69504) according to the manufacturer’s instructions. The measurement of cytosolic mitochondrial DNA (mtDNA) via qPCR is determined by the ratio of the cytosolic fraction to the whole-cell DNA fraction.

The following primers were used:

hMT-Dloop-F: CATCTGGTTCCTACTTCAGGG

hMT-Dloop-R: CCGTGAGTGGTTAATAGGGTG

### RNA-seq

RNA-seq was performed as described previously ^40^. RNA quality was checked using Agilent Tapestation. RNA-seq libraries were prepared from total RNA using polyA selection followed by the NEBNext Ultra II Directional RNA library kit workflow (New England Biolabs). Sequencing was performed on the Illumina HiSeq 2500 instrument, resulting in approximately 30 million 50 bp reads per sample. Sequencing reads were mapped in a splice-aware fashion to the human reference transcriptome (hg19 assembly) using STAR ^43^. Read counts over transcripts were calculated using HTSeq^44^ based on the Ensembl annotation for GRCh38/hg19 assembly. For the differential expression analysis, we used the EdgeR method ^44,45^ and defined differentially expressed genes (DEGs) based on the cutoffs of 2-fold change in expression value and false discovery rates (FDR) below 0.05. GO analysis was performed using EnrichR^46^. For heatmap visualization of SASP genes, the SASP factors curated previously were used as a reference^47^. Proliferating controls were described previously ^40^ and heatmaps were generated using an online tool^48^.

### Statistical analyses

Unpaired two-tailed Student’s t-test was used for comparison between two groups. One-way ANOVA coupled with Tukey’s post hoc test was used for comparisons over two groups. Significance was considered when p value was less than 0.05.

### Data availability

RNA-seq data have been deposited in the NCBI Gene Expression Omnibus (GEO) database under accession number GSE272306. Other original data are available upon reasonable request.

## Extended Data Figure Legends

**Extended Data Figure 1.**
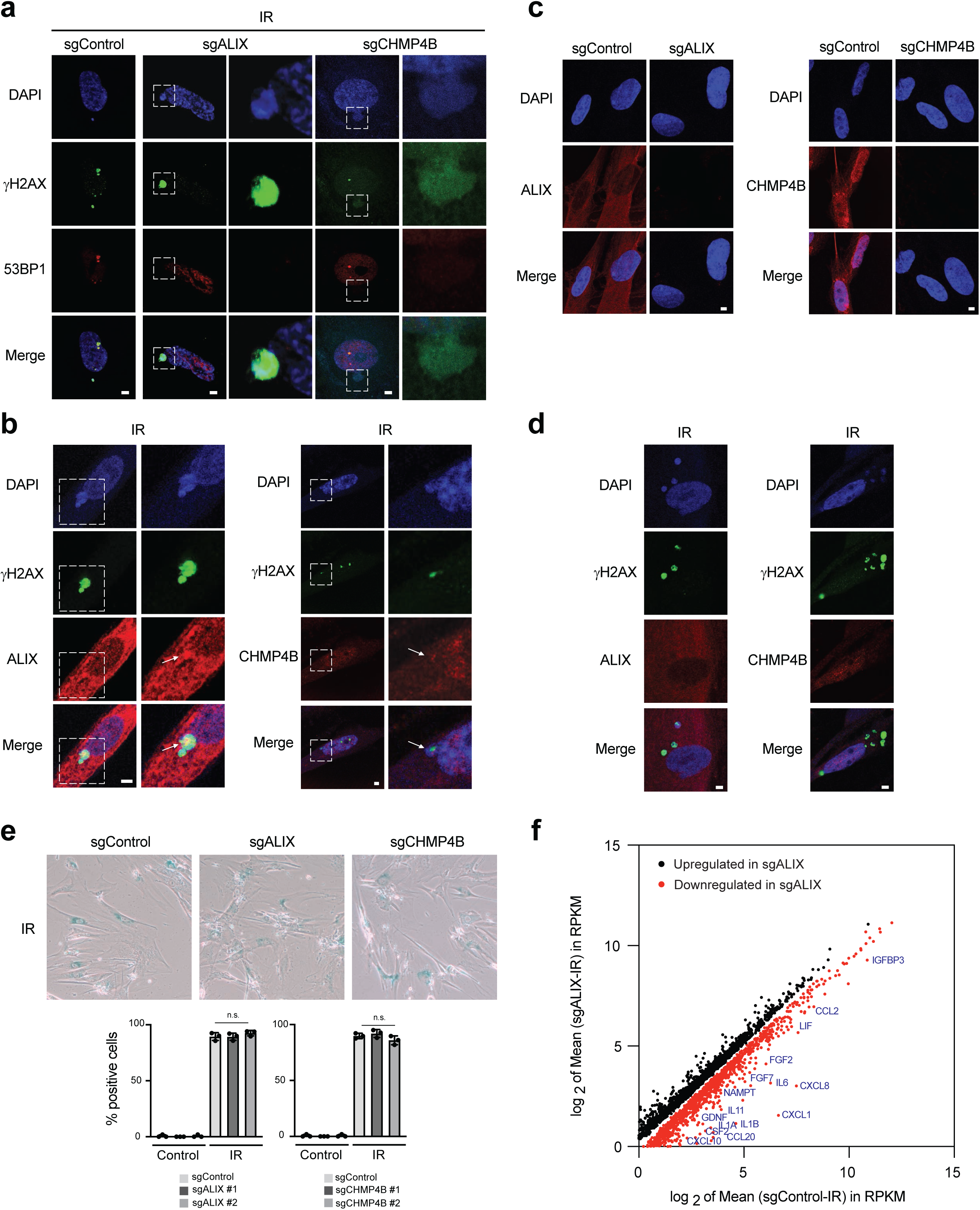
Roles of ESCRT-III in mediating CCF nuclear egress. **a**, Related to Fig. 1b, additional images of CCF and nuclear membrane blebs in control, CHMP4B, or ALIX deficient senescent IMR90 induced by IR. Scale bar: 1 μm. **b**, Related to Fig. 1e, additional images of CHMP4B and ALIX localization at the nuclear membrane blebs of senescent cells. Scale bar: 1 μm. **c**, Related to Fig. 1e, antibody validation using KO cells. CHMP4B or ALIX-deficient proliferating IMR90 cells were stained with CHMP4B or ALIX antibody and imaged under a confocal microscopy. Note that the signal is lost in CRISPR KO cells. Scale bar: 1 μm.**d**, CHMP4B and ALIX are not enriched at CCFs of senescent cells. Note that the cytoplasmic CCFs stain negative for CHMP4B and ALIX. Scale bar: 1 μm. **e**, SA-β-gal analyses of ESCRT-III-deficient cells. IMR90 cells were left untreated or induced to senescence by IR and fixed after 14 days. The cells were stained with SA-β-gal kit and representative images are presented (left). (Right) Quantification of SA-β-gal positive cells. Results shown are the mean values with s.d. from four randomly selected fields with over 200 cells. P values are calculated with one-way ANOVA coupled with Tukey’s post hoc test. n.s.: non-significant. **f**, Related to Fig. 1i and 1j, RNA-seq analyses of sgControl and sgALIX senescent cells. The mean values of RPKM of gene expression were plotted and key SASP genes were annotated. Note that sgALIX senescent cells show reduced expression of multiple SASP genes.

**Extended Data Figure 2.**
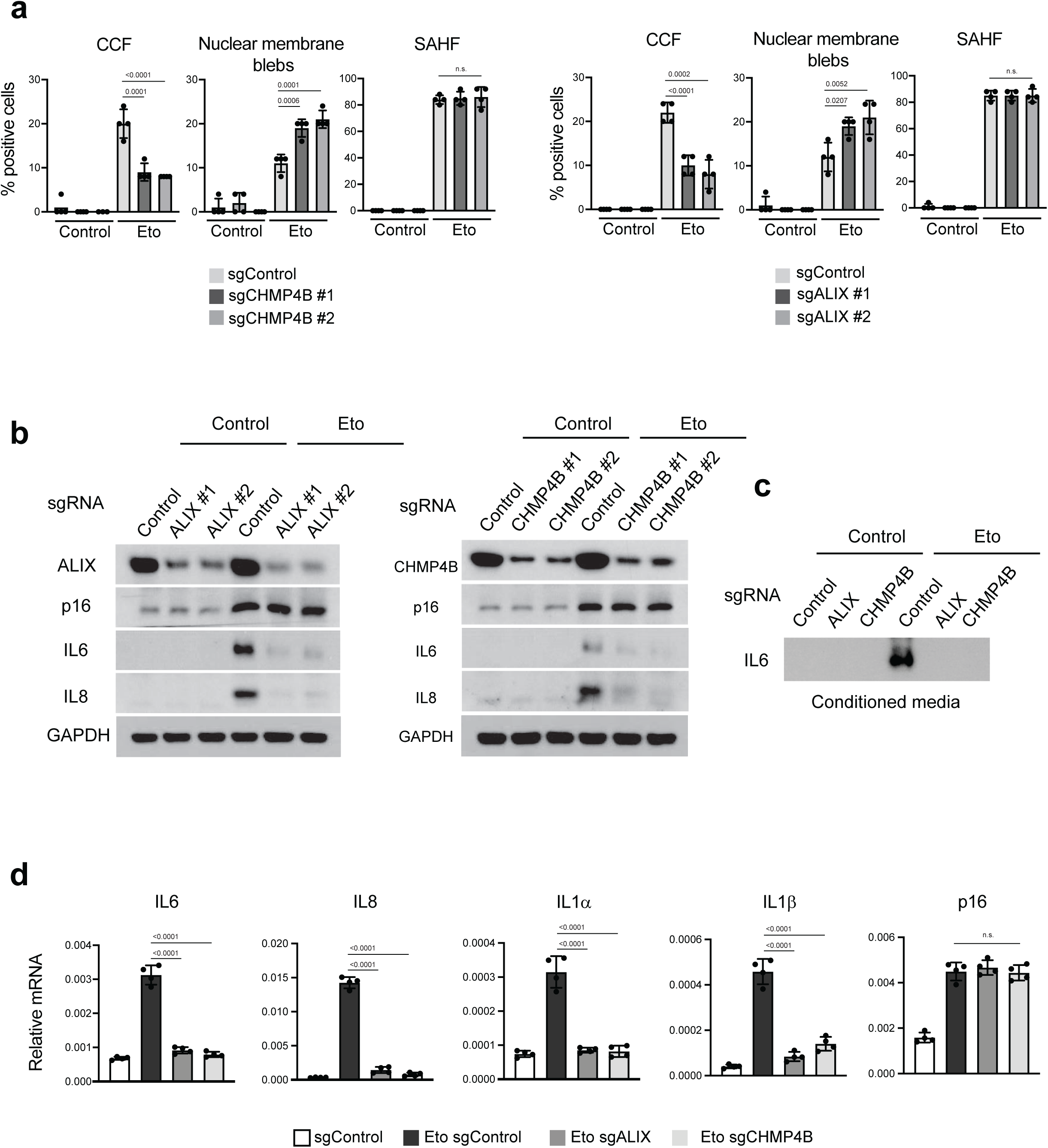
ESCRT-III is required for CCF and SASP in etoposide-induced senescent IMR90 cells. **a**, IMR90 cells were stably infected with lentivirus encoding sgControl, sgCHMP4B, or sgALIX. The cells were treated with 50 uM etoposide for 48 hours and were harvested on day 12. The cells were then fixed and subjected to imaging analyses and quantification. Results shown are the mean values with s.d. from four randomly selected fields with over 200 cells. P values are calculated with one-way ANOVA coupled with Tukey’s post hoc test. n.s.: non-significant. **b**, Cells as in **a** were analyzed by immunoblotting. **c**, The conditioned media were analyzed by IL6 immunoblotting. The amounts of media loaded were normalized to total cellular proteins. See Materials and Methods for details. **d**, Control or etoposide-induced senescent IMR90 cells were analyzed by RT-qPCR for indicated genes. Results shown are the mean values with s.d., normalized to those of Lamin A/C. P values are calculated with one-way ANOVA coupled with Tukey’s post hoc test. n.s.: non-significant.

**Extended Data Figure 3.**
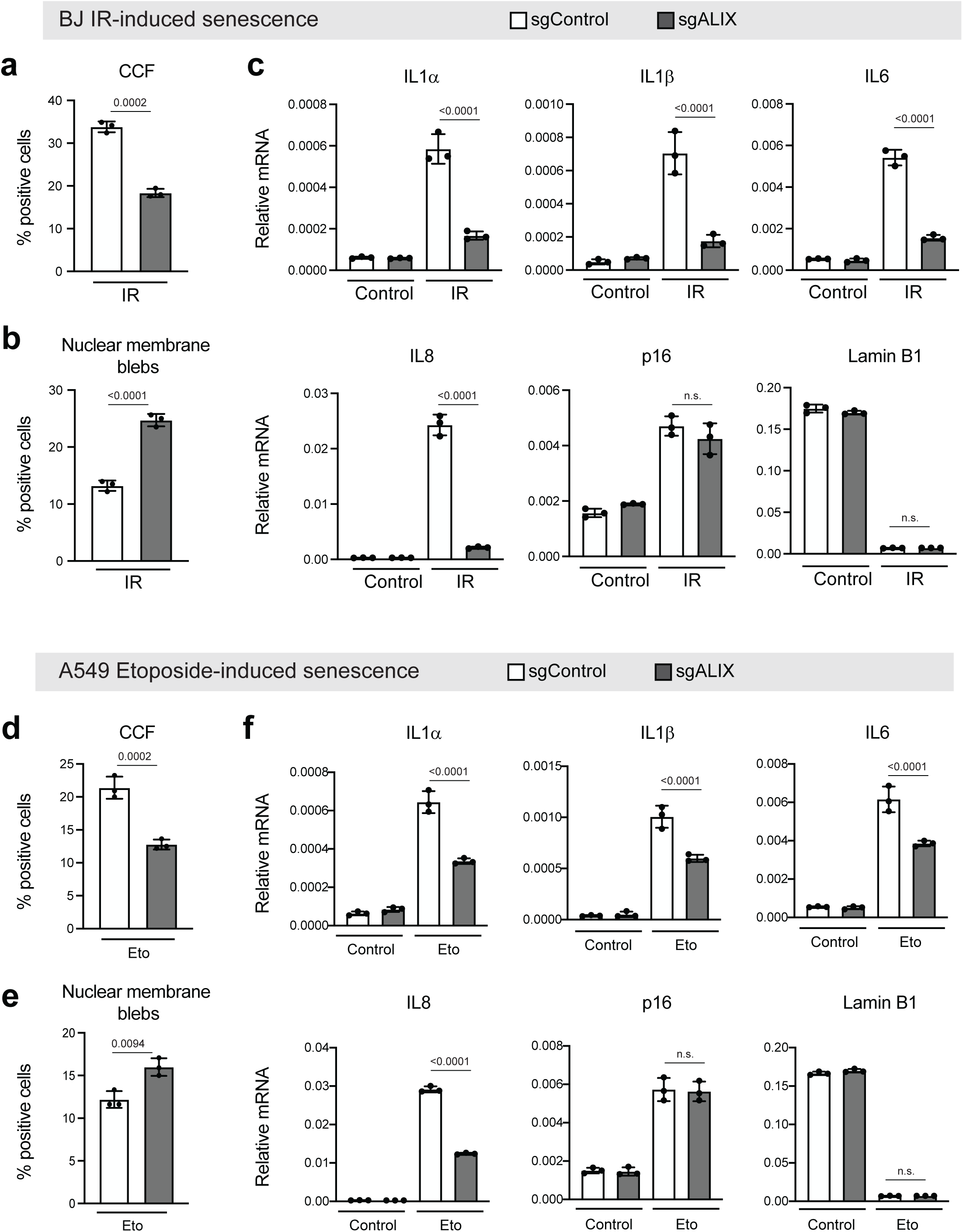
ALIX is required for CCF and the SASP in senescence of BJ fibroblasts and A549 cells. **a and b**, Primary BJ fibroblasts were stably infected with CRISPR constructs encoding sgControl or sgALIX. The cells were then induced to senescence with 20 Gy of IR and harvested on day 12, fixed, and subjected to imaging analyses and quantification. Bar graphs show mean values with s.d. from four randomly selected fields with over 200 cells. P values are calculated with one-way ANOVA coupled with Tukey’s post hoc test. **c**, The BJ cells were analyzed by RT-qPCR analyses for indicated genes. Results shown are mean values with s.d., normalized to those of Lamin A/C. P values are calculated with one-way ANOVA coupled with Tukey’s post hoc test. n.s.: non-significant. **d and e**, A549 cells were stably infected with CRISPR constructs encoding sgControl or sgALIX. The cells were then induced to senescence with 5 μM etoposide and harvested on day 14, fixed, and subjected to imaging analyses and quantification. Bar graphs show mean values with s.d. from three randomly selected fields with over 200 cells. P values are calculated with one-way ANOVA coupled with Tukey’s post hoc test. **f**, The A549 cells were analyzed by RT-qPCR analyses for indicated genes. Results shown are mean values with s.d., normalized to those of Lamin A/C. P values are calculated with one-way ANOVA coupled with Tukey’s post hoc test. n.s.: non-significant.

**Extended Data Figure 4.**
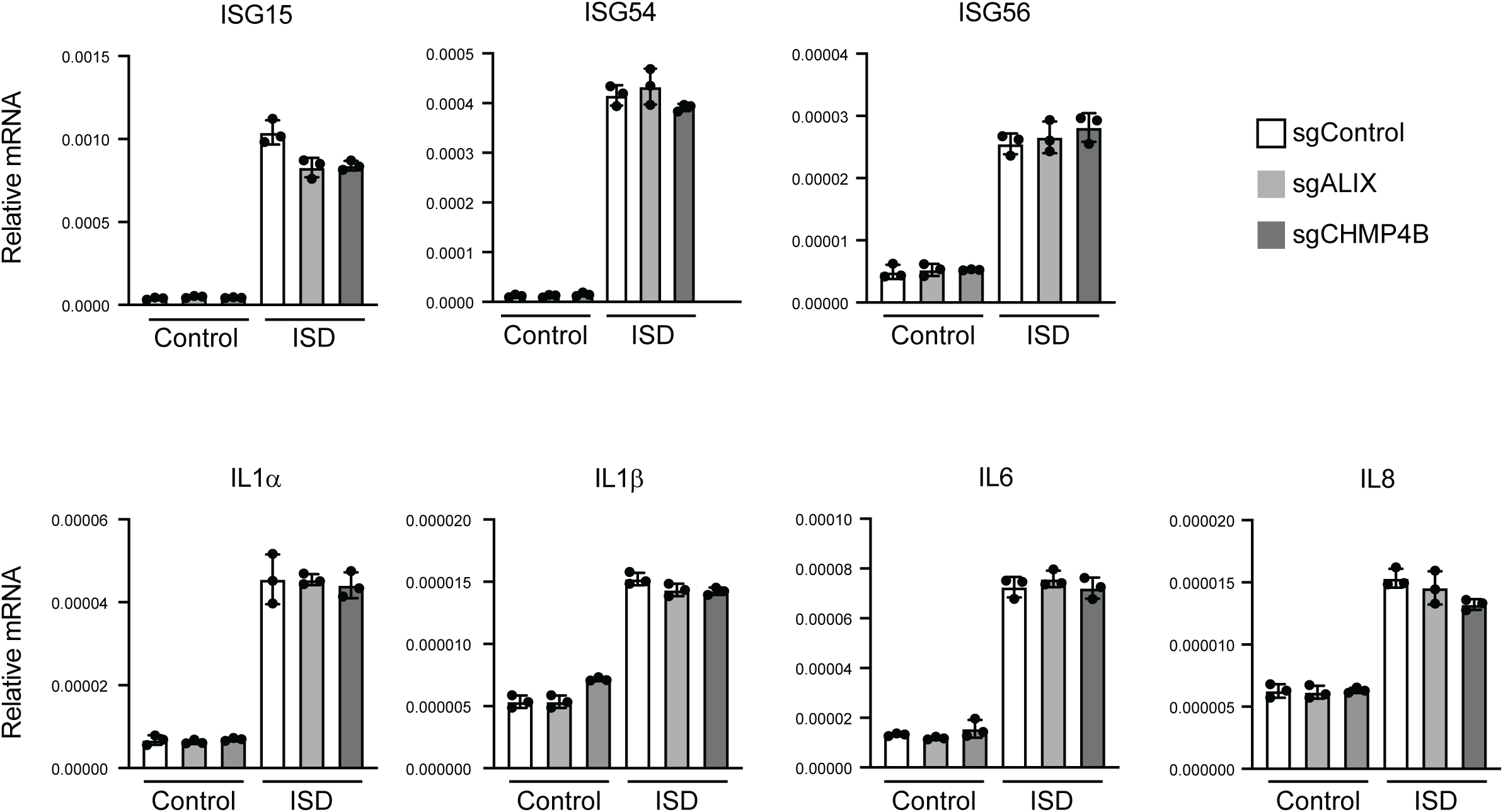
ESCRT-III has minimal roles in acute dsDNA-induced inflammatory responses. IMR90 cells were stably infected with CRISPR constructs encoding sgControl, sgCHMP4B, or sgALIX. The cells were then transfected with ISD and were harvested 24 hours later, followed by RT-qPCR analyses with indicated genes. Results shown are the mean values with s.d., normalized to those of Lamin A/C.

**Extended Data Figure 5.**
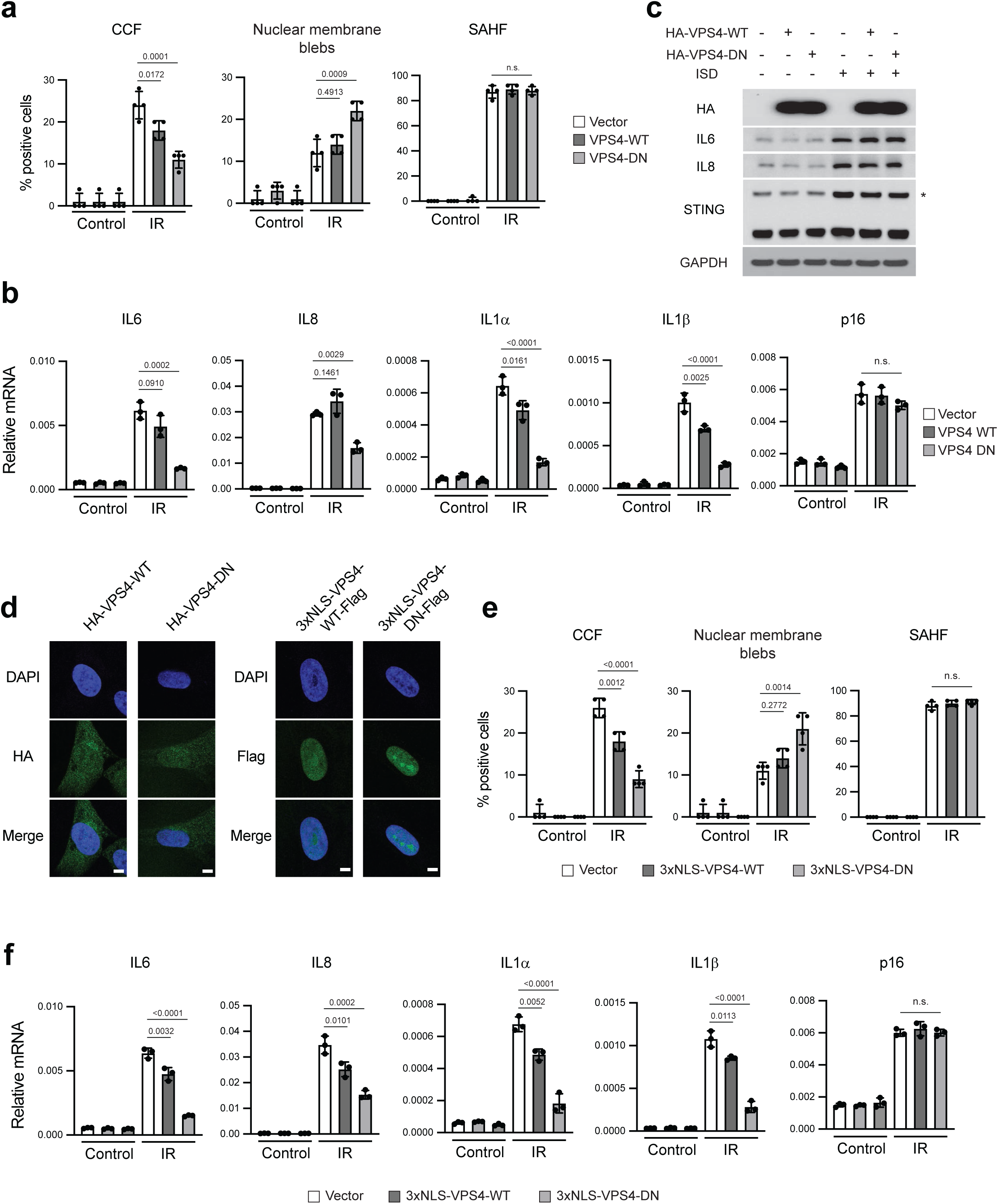
VPS4, and specifically nuclear VPS4, regulates CCF nuclear egress and the SASP. **a-c**, IMR90 cells were stably infected with constructs encoding a vector, HA-VPS4-WT, or HA-VPS4-DN. The cells were then induced to senescence by IR and harvested at day 14 (**a** and **b**). The cells were fixed and subjected to imaging analyses (**a**). Bar graphs show mean values with s.d. from four randomly selected fields with over 200 cells. P values are calculated with one-way ANOVA coupled with Tukey’s post hoc test. n.s.: non-significant. **b**, The cells were analyzed by RT-qPCR for indicated genes. Results shown are mean values with s.d., normalized to those of Lamin A/C. P values are calculated with one-way ANOVA coupled with Tukey’s post hoc test. n.s.: non-significant. **c**, Cells were transfected with ISD and were harvested 24 hours later. The lysates were analyzed by immunoblotting. STING western was performed under non-reducing conditions. * denotes STING dimer. **d-f,** IMR90 cells were stably infected with constructs encoding a vector, 3xNLS-VPS4-WT-Flag, or 3xNLS-VPS4-DN-Flag. The cells were imaged under a confocal microscopy (**d**). Scale bar: 5 μm. **e and f**, The cells as indicated were induced to senescence by IR and harvested at day 14. The cells were fixed and subjected to imaging analyses (**e**). Bar graphs show mean values with s.d. from four randomly selected fields with over 200 cells. P values are calculated with one-way ANOVA coupled with Tukey’s post hoc test. n.s.: non-significant. **f**, The cells were analyzed by RT-qPCR analyses with indicated genes. Results shown are mean values with s.d., normalized to those of Lamin A/C. P values are calculated with one-way ANOVA coupled with Tukey’s post hoc test. n.s.: non-significant.

**Extended Data Figure 6.**
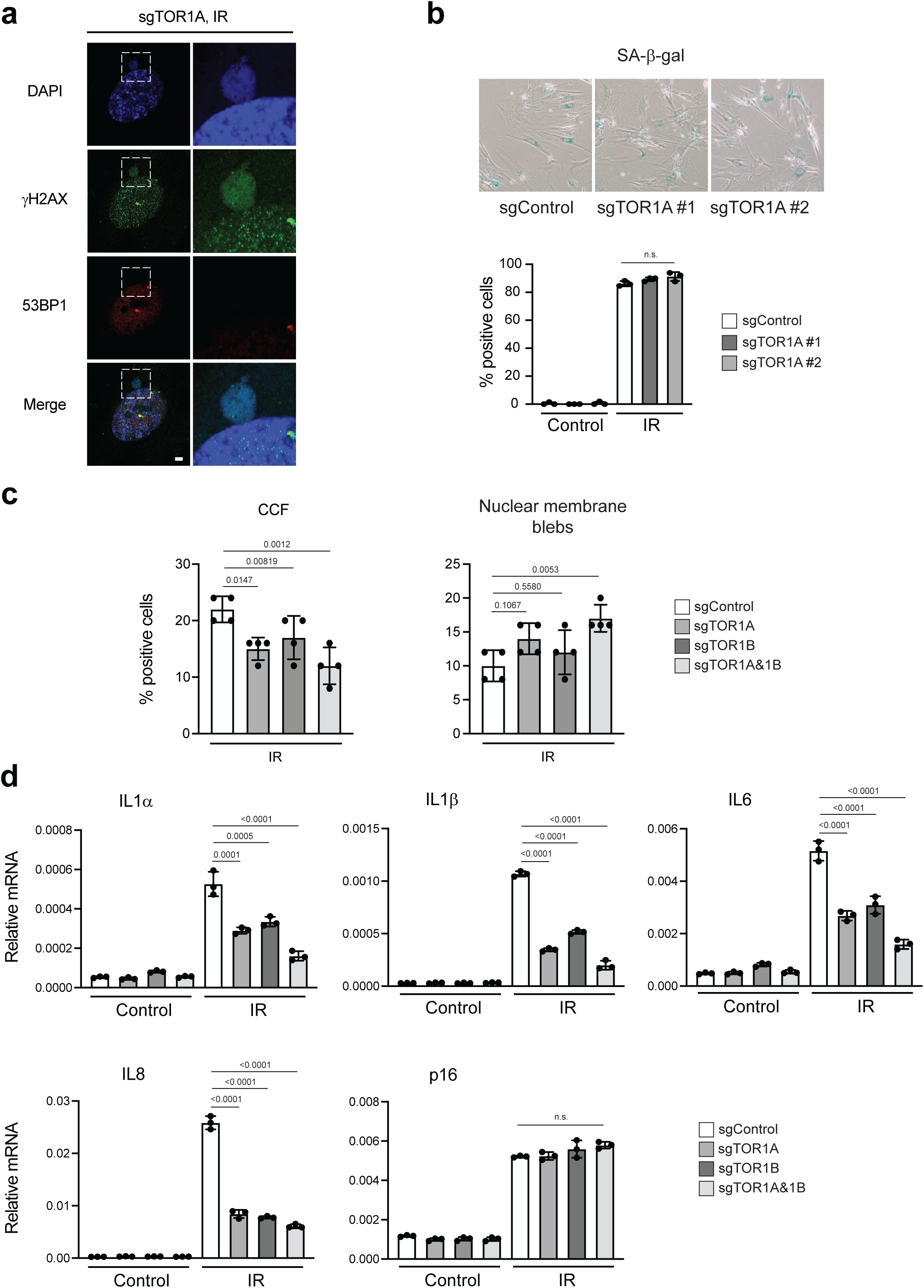
Torsin complex is required for CCF nuclear egress and the SASP. **a**, Related to Fig. 2a, additional images of CCF and nuclear membrane blebs in TOR1A-deficient senescent IMR90 cells. Scale bar: 5 μm. **b**, SA-β-gal analyses of TOR1A-deficient cells. IMR90 cells were left untreated or induced to senescence by IR and fixed after 14 days. The cells were stained with SA-β-gal kit and representative images are presented (top). (Bottom) Quantification of SA-β-gal positive cells. Results shown are the mean values with s.d. from four randomly selected fields with over 200 cells. P values are calculated with one-way ANOVA coupled with Tukey’s post hoc test. n.s.: non-significant. **c and d**, IMR90 cells were stably infected with CRISPR constructs encoding sgControl, sgTOR1A, sgTOR1B, or sgTOR1A plus sgTOR1B. The cells were induced to senescence by IR and analyzed by imaging (**c**). Bar graphs show mean values with s.d.. from four randomly selected fields with over 200 cells. P values are calculated with one-way ANOVA coupled with Tukey’s post hoc test. **d**, RT-qPCR analyses with indicated genes. Bar graphs show mean values with s.d.. Results are normalized to those of Lamin A/C. P values are calculated with one-way ANOVA coupled with Tukey’s post hoc test. n.s.: non-significant.

**Extended Data Figure 7.**
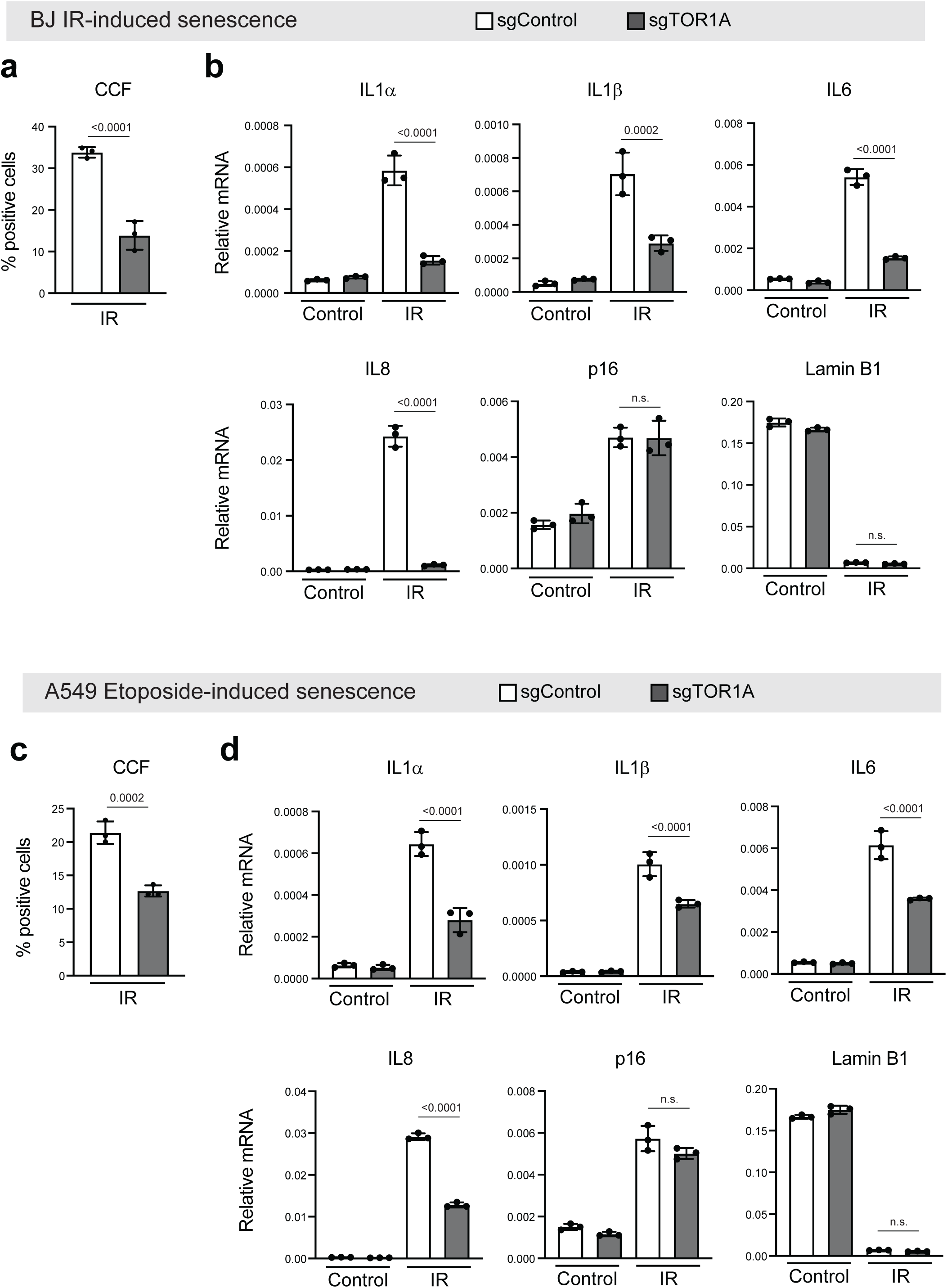
TOR1A is required for CCF and the SASP in senescence of BJ fibroblasts and A549 cells. **a**, Primary BJ fibroblasts were stably infected with CRISPR constructs encoding sgControl or sgTOR1A. The cells were then induced to senescence with 20 Gy of IR and harvested on day 12, fixed, and subjected to imaging analyses and quantification. Bar graphs show mean values with s.d. from three randomly selected fields with over 200 cells. **b**, Cells as in **a** were analyzed by RT-qPCR analyses for indicated genes. Results shown are mean values with s.d., normalized to those of Lamin A/C. **c**, A549 cells were stably infected with CRISPR constructs encoding sgControl or sgTOR1A. The cells were then induced to senescence with 5μM of etoposide and harvested on day 14, fixed, and subjected to imaging analyses and quantification. Bar graphs show mean values with s.d. from three randomly selected fields with over 200 cells. **d**, Cells as in **c** were analyzed by RT-qPCR analyses for indicated genes. Results shown are mean values with s.d., normalized to those of Lamin A/C. P values in this figure were calculated with one-way ANOVA coupled with Tukey’s post hoc test. The same sgControl samples were used from Extended Data Figure 3. n.s.: non-significant.

**Extended Data Figure 8.**
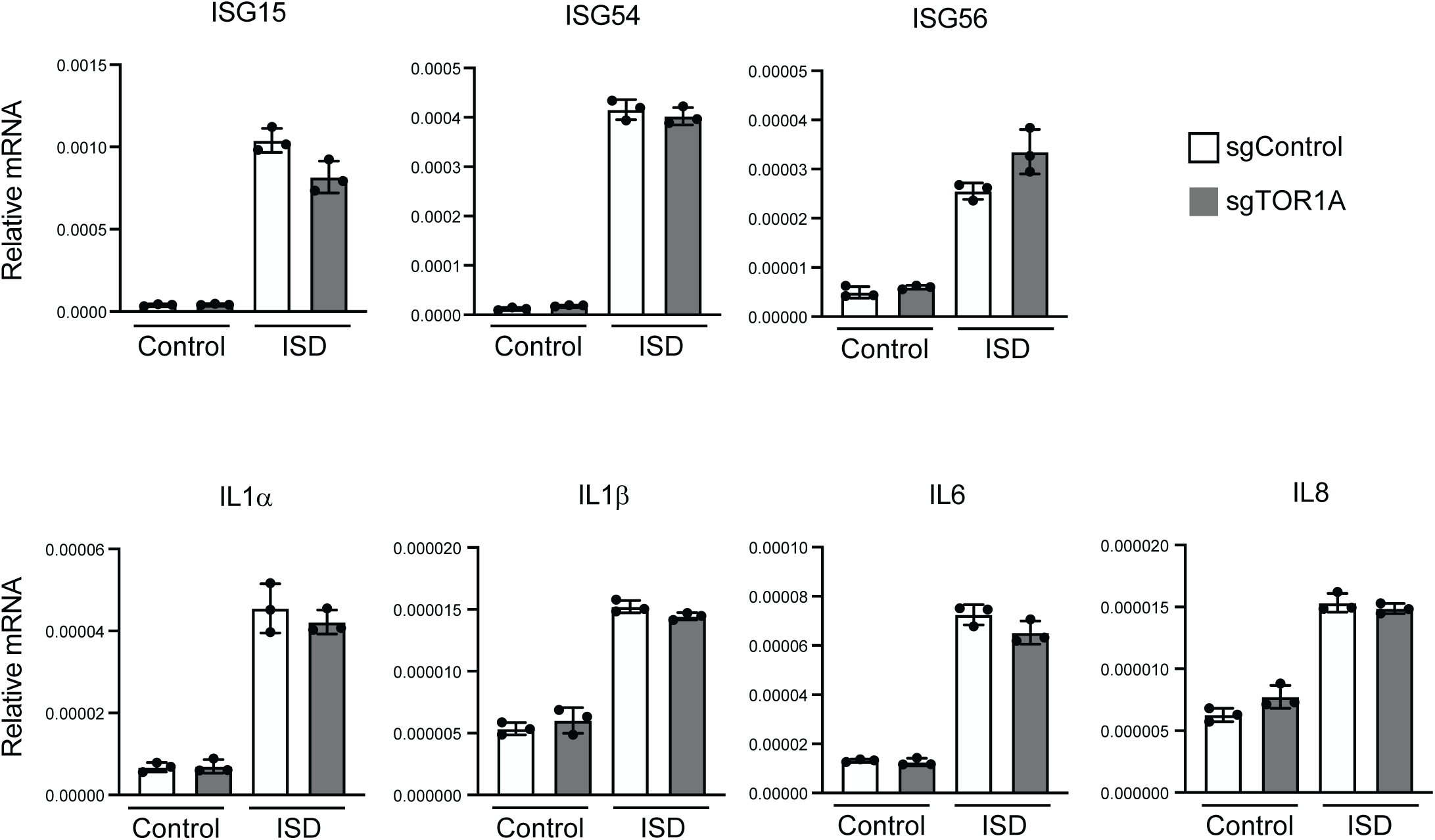
TOR1A has minimal roles in acute dsDNA-induced inflammatory responses. IMR90 cells were stably infected with CRISPR constructs encoding sgControl or sgTOR1A. The cells were then transfected with ISD and were harvested 24 hours later, followed by RT-qPCR analyses with indicated genes. Results shown are mean values with s.d., normalized to those of Lamin A/C.

**Extended Data Figure 9.**
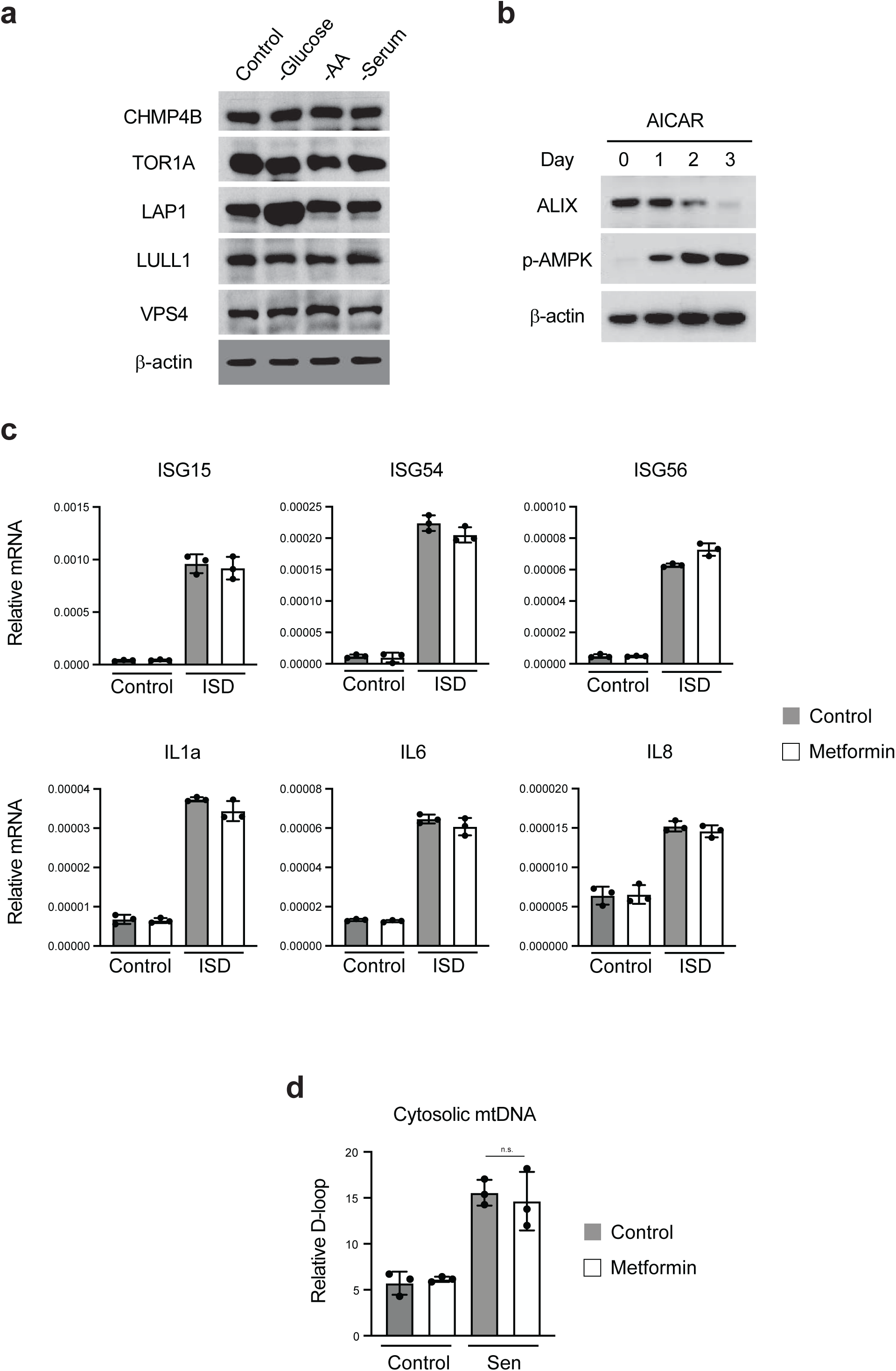
Effects of glucose limitation and metformin on nuclear egress proteins and cytosolic DNA sensing pathway. **a**, Related to Fig. 3a, western blotting of proteins involved in nuclear egress upon glucose starvation. **b**, Related to Fig. 3g, IMR90 cells were cultured in 5 mM glucose media in the presence of 1 mM AICAR for indicated days, and were analyzed by immunoblotting. **c**, IMR90 cells were cultured in 5 mM media with or without 5 mM metformin for 2 days. The cells were then transfected with ISD and were harvested 24 hours later, followed by RT-qPCR analyses with indicated genes. Results shown are mean values with s.d., normalized to those of Lamin A/C. **d**, IMR90 cells were induced to senescence by IR. On day 8, the cells were cultured in 5 mM glucose media with or without 5 mM metformin. The cells were then harvested on day 14 and analyzed for cytosolic mitochondrial DNA using mitochondrial D-loop primer. See Methods for more details. Results shown are mean values with s.d.. n.s.: non-significant, as calculated with one-way ANOVA coupled with Tukey’s post hoc test.

**Extended Data Figure 10.**
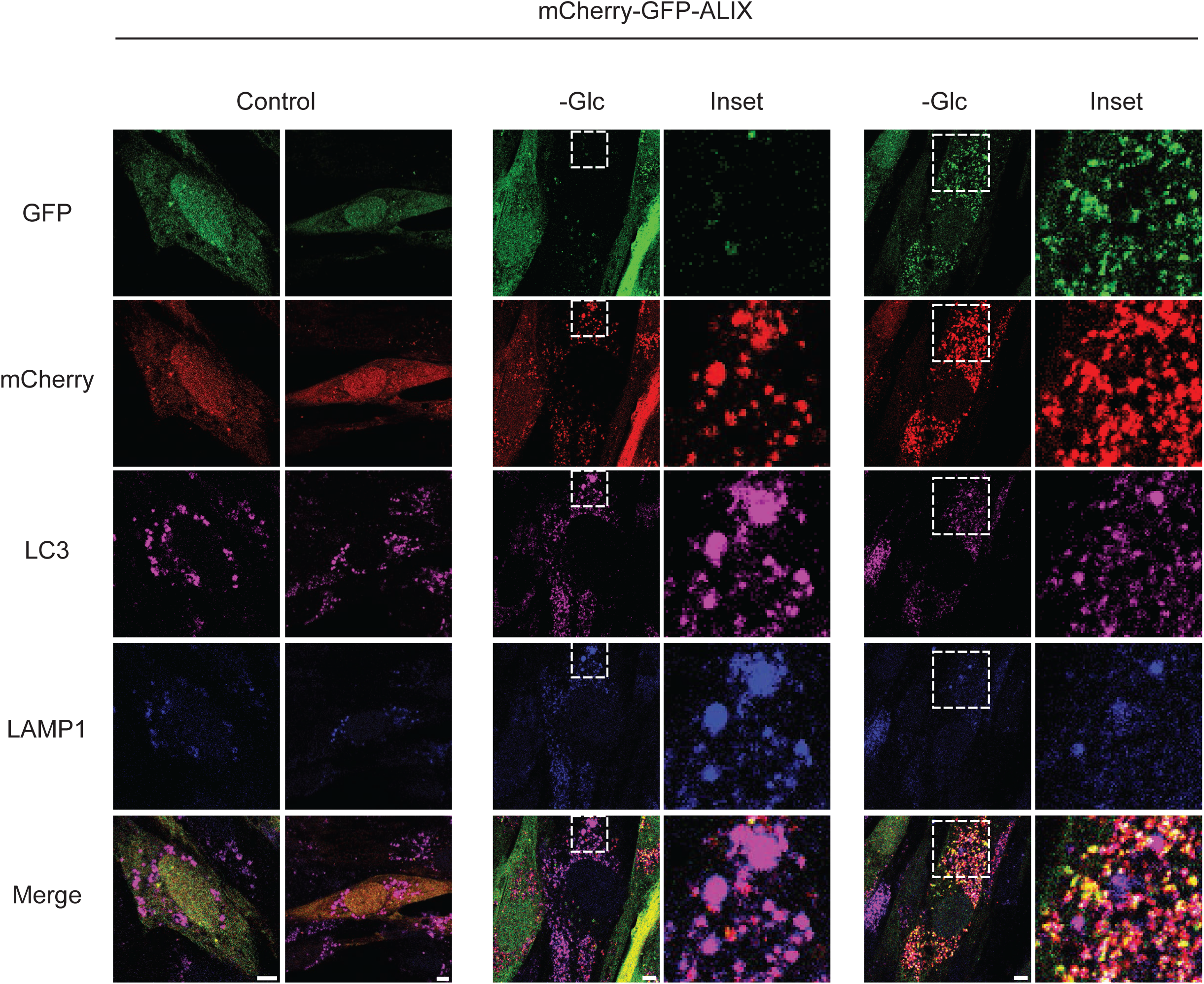
ALIX undergoes autophagy-lysosome degradation upon glucose limitation. mCherry-GFP-ALIX was stably expressed in IMR90 cells. The cells were left untreated or glucose starved for 1 day and were subjected to immunostaining and imaging under a confocal microscopy. Scale bar: 5 μm.

## References

1. Franceschi, C. & Campisi, J. Chronic inflammation (inflammaging) and its potential contribution to age-associated diseases. J. Gerontol. A Biol. Sci. Med. Sci. 69 **Suppl 1**, S4–9 (2014).

2. López-Otín, C., Blasco, M. A., Partridge, L., Serrano, M. & Kroemer, G. Hallmarks of aging: An expanding universe. Cell 186, 243–278 (2023).

3. Miller, K. N. et al. Cytoplasmic DNA: sources, sensing, and role in aging and disease. Cell 184, 5506–5526 (2021).

4. Ivanov, A. et al. Lysosome-mediated processing of chromatin in senescence. J. Cell Biol. 202, 129–143 (2013).

5. Dou, Z. et al. Cytoplasmic chromatin triggers inflammation in senescence and cancer. Nature 550, 402–406 (2017).

6. Vizioli, M. G. et al. Mitochondria-to-nucleus retrograde signaling drives formation of cytoplasmic chromatin and inflammation in senescence. Genes Dev. 34, 428–445 (2020).

7. Yang, H., Wang, H., Ren, J., Chen, Q. & Chen, Z. J. cGAS is essential for cellular senescence. Proc. Natl. Acad. Sci. U. S. A. 114, E4612–E4620 (2017).

8. Glück, S. et al. Innate immune sensing of cytosolic chromatin fragments through cGAS promotes senescence. Nat. Cell Biol. 19, 1061–1070 (2017).

9. Gulen, M. F. et al. cGAS–STING drives ageing-related inflammation and neurodegeneration. Nature 620, 374–380 (2023).

10. Dou, Z. et al. Autophagy mediates degradation of nuclear lamina. Nature 527, 105–109 (2015).

11. Klupp, B. G. & Mettenleiter, T. C. The Knowns and Unknowns of Herpesvirus Nuclear Egress. Annu Rev Virol 10, 305–323 (2023).

12. Arii, J. et al. ESCRT-III mediates budding across the inner nuclear membrane and regulates its integrity. Nat. Commun. 9, 3379 (2018).

13. Arii, J. Host and Viral Factors Involved in Nuclear Egress of Herpes Simplex Virus 1. Viruses 13, (2021).

14. Maric, M. et al. A functional role for TorsinA in herpes simplex virus 1 nuclear egress. J. Virol. 85, 9667–9679 (2011).

15. Jokhi, V. et al. Torsin mediates primary envelopment of large ribonucleoprotein granules at the nuclear envelope. Cell Rep. 3, 988–995 (2013).

16. Narita, M. et al. Rb-mediated heterochromatin formation and silencing of E2F target genes during cellular senescence. Cell 113, 703–716 (2003).

17. McCullough, J., Frost, A. & Sundquist, W. I. Structures, Functions, and Dynamics of ESCRT-III/Vps4 Membrane Remodeling and Fission Complexes. Annu. Rev. Cell Dev. Biol. 34, 85–109 (2018).

18. Hurley, J. H. ESCRTs are everywhere. EMBO J. 34, 2398–2407 (2015).

19. Vietri, M., Radulovic, M. & Stenmark, H. The many functions of ESCRTs. Nat. Rev. Mol. Cell Biol. 21, 25–42 (2020).

20. Laudermilch, E. & Schlieker, C. Torsin ATPases: structural insights and functional perspectives. Curr. Opin. Cell Biol. 40, 1–7 (2016).

21. Zhao, C., Brown, R. S. H., Chase, A. R., Eisele, M. R. & Schlieker, C. Regulation of Torsin ATPases by LAP1 and LULL1. Proc. Natl. Acad. Sci. U. S. A. 110, E1545–54 (2013).

22. Sosa, B. A. et al. How lamina-associated polypeptide 1 (LAP1) activates Torsin. Elife 3, e03239 (2014).

23. Goodchild, R. E. & Dauer, W. T. The AAA+ protein torsinA interacts with a conserved domain present in LAP1 and a novel ER protein. J. Cell Biol. 168, 855–862 (2005).

24. Finkel, T. The metabolic regulation of aging. Nat. Med. 21, 1416–1423 (2015).

25. Herzig, S. & Shaw, R. J. AMPK: guardian of metabolism and mitochondrial homeostasis. Nat. Rev. Mol. Cell Biol. 19, 121–135 (2018).

26. Gwinn, D. M. et al. AMPK phosphorylation of raptor mediates a metabolic checkpoint. Mol. Cell 30, 214–226 (2008).

27. Victorelli, S. et al. Apoptotic stress causes mtDNA release during senescence and drives the SASP. Nature 622, 627–636 (2023).

28. Mizushima, N., Yoshimori, T. & Levine, B. Methods in mammalian autophagy research. Cell 140, 313–326 (2010).

29. Lamark, T. & Johansen, T. Mechanisms of Selective Autophagy. Annu. Rev. Cell Dev. Biol. 37, 143–169 (2021).

30. Kulkarni, A. S., Gubbi, S. & Barzilai, N. Benefits of Metformin in Attenuating the Hallmarks of Aging. Cell Metab. 32, 15–30 (2020).

31. Foretz, M., Guigas, B., Bertrand, L., Pollak, M. & Viollet, B. Metformin: from mechanisms of action to therapies. Cell Metab. 20, 953–966 (2014).

32. Martin-Montalvo, A. et al. Metformin improves healthspan and lifespan in mice. Nat. Commun. 4, 2192 (2013).

33. Yang, Y. et al. Metformin decelerates aging clock in male monkeys. Cell 187, 6358–6378.e29 (2024).

34. Tobar, N. et al. Metformin acts in the gut and induces gut-liver crosstalk. Proc. Natl. Acad. Sci. U. S. A. 120, e2211933120 (2023).

35. McCreight, L. J., Bailey, C. J. & Pearson, E. R. Metformin and the gastrointestinal tract. Diabetologia 59, 426–435 (2016).

36. Moiseeva, O. et al. Metformin inhibits the senescence-associated secretory phenotype by interfering with IKK/NF-κB activation. Aging Cell 12, 489–498 (2013).

37. van Vliet, T. et al. Physiological hypoxia restrains the senescence-associated secretory phenotype via AMPK-mediated mTOR suppression. Mol. Cell 81, 2041–2052.e6 (2021).

38. Stowe, R. P. et al. Chronic herpesvirus reactivation occurs in aging. Exp. Gerontol. 42, 563–570 (2007).

39. Bennett, J. M. et al. Inflammation and reactivation of latent herpesviruses in older adults. Brain Behav. Immun. 26, 739–746 (2012).

40. Wang, Y. et al. Nuclear autophagy interactome unveils WSTF as a constitutive nuclear inhibitor of inflammation. bioRxiv 2022.10.04.510822 (2022) doi:10.1101/2022.10.04.510822.

41. Xu, C. et al. SIRT1 is downregulated by autophagy in senescence and ageing. Nat. Cell Biol. 22, 1170–1179 (2020).

42. Onorati, A. et al. Upregulation of PD-L1 in Senescence and Aging. Mol. Cell. Biol. 42, e0017122 (2022).

43. Dobin, A. et al. STAR: ultrafast universal RNA-seq aligner. Bioinformatics 29, 15–21 (2013).

44. Anders, S., Pyl, P. T. & Huber, W. HTSeq--a Python framework to work with high-throughput sequencing data. Bioinformatics 31, 166–169 (2015).

45. Robinson, M. D., McCarthy, D. J. & Smyth, G. K. edgeR: a Bioconductor package for differential expression analysis of digital gene expression data. Bioinformatics 26, 139–140 (2010).

46. Kuleshov, M. V. et al. Enrichr: a comprehensive gene set enrichment analysis web server 2016 update. Nucleic Acids Res. 44, W90–7 (2016).

47. Freund, A., Orjalo, A. V., Desprez, P.-Y. & Campisi, J. Inflammatory networks during cellular senescence: causes and consequences. Trends Mol. Med. 16, 238–246 (2010).

48. Babicki, S. et al. Heatmapper: web-enabled heat mapping for all. Nucleic Acids Res. 44, W147–53 (2016).

